# *Actinobacillus pleuropneumoniae* encodes multiple phase-variable DNA methyltransferases that comprise distinct phasevarions

**DOI:** 10.1101/2022.11.17.516983

**Authors:** Nusrat Nahar, Greg Tram, Freda E-C Jen, Zachary N. Phillips, Lucy A. Weinert, Janine T. Bossé, Jafar S. Jabbari, Quentin Gouil, Mei R. M. Du, Matthew E. Ritchie, Rory Bowden, Paul R. Langford, Alexander W. Tucker, Michael P. Jennings, Conny Turni, Patrick J. Blackall, John M. Atack

**Affiliations:** Institute for Glycomics, Griffith University, Gold Coast, Queensland, 4222, Australia; Department of Veterinary Medicine, University of Cambridge, Cambridge CB3 0ES, UK.; Section of Paediatric Infectious Disease, Imperial College London, St Mary’s Campus, London, W2 1PG, United Kingdom; Walter and Eliza Hall Institute of Medical Research, Parkville, Victoria, 3052, Australia; Department of Medical Biology, The University of Melbourne, Parkville, 3010, Australia; Queensland Alliance for Agriculture and Food Innovation, The University of Queensland, St. Lucia, Queensland, 4072, Australia; School of Environment and Science, Griffith University, Gold Coast, Queensland, 4222, Australia

## Abstract

*Actinobacillus pleuropneumoniae* is the cause of porcine pleuropneumonia, a severe respiratory tract infection that is responsible for major economic losses to the swine industry. Many host-adapted bacterial pathogens encode systems known as phasevarions (phase-variable regulons). Phasevarions result from variable expression of cytoplasmic DNA methyltransferases. Variable expression results in genome-wide methylation differences within a bacterial population, leading to altered expression of multiple genes via epigenetic mechanisms. Our examination of a diverse population of *A. pleuropneumoniae* strains determined that Type I and Type III DNA methyltransferases with the hallmarks of phase variation were present in this species. We demonstrate that phase variation is occurring in these methyltransferase, and show associations between particular Type III methyltransferase alleles and serovar. Using Pacific BioSciences Single-Molecule, Real-Time (SMRT) sequencing and Oxford Nanopore sequencing, we demonstrate the presence of the first ever characterised phase-variable, cytosine-specific Type III DNA methyltransferase. Phase variation of distinct Type III DNA methyltransferase variants results in the regulation of distinct phasevarions, and in multiple phenotypic differences relevant to pathobiology. Our characterisation of these newly described phasevarions in *A. pleuropneumoniae* will aid in the selection of stably expressed antigens, and direct and inform development of a rationally designed subunit vaccine against this major veterinary pathogen.

## INTRODUCTION

Porcine pleuropneumonia, a severe respiratory disease of pigs, is caused by *Actinobacillus pleuropneumoniae* and is associated with major annual global economic losses to the swine industry (1). As *A. pleuropneumoniae* is an obligate coloniser of the porcine respiratory tract, the organism is usually isolated from lung and tonsils of acutely or chronically infected pigs (2–4). *A. pleuropneumoniae* is transmitted by direct contact with infected pigs or aerosol droplets (5). On the basis of distinct capsular polysaccharides, 19 serovars have been identified (6). Available vaccines reduce the severity of disease without impacting colonisation, but these are often serovar specific; no vaccine exists to protect against all strains, and neither antibiotics nor current vaccines eliminate the spread of bacteria (7, 8).

Phase variation is the rapid and reversible switching of gene expression, and is typically associated with bacterial virulence determinants (9–12), particularly genes that encode surface structures such as adhesins (13, 14), pili (15), capsule (16) and lipooligosaccharide (17). Phase variation serves as an extra contingency strategy for a bacterial population by generating phenotypic diversity. The two most common mechanisms of phase variation are i) slipped strand mispairing of simple DNA sequence repeats (SSRs) located within an open reading frame of a gene, and ii) sequence variation through homologous recombination between expressed and silent loci that contain inverted repeats (IRs) (18–21). Variation in length of locus located SSRs results in a biphasic ON-OFF switching of gene expression. This is due to changes in the reading frame commensurate with the number of individual repeats present in the SSR tract. Homologous recombination between expressed and silent loci via IRs, also referred to as antigenic variation, means that a protein is always expressed, but as multiple allelic variants within a population dependent on the sequence of the encoding gene that is expressed. Phase variation can complicate vaccine development as it results in an unstable antigenic repertoire, which can allow, for example, escape of a vaccine primed immune response.

Recent surveys of the restriction enzyme database REBASE (22) have demonstrated that almost 20% of Type I and Type III DNA methyltransferases contain the hallmarks of phase-variable expression (SSR tracts or multiple variable loci that contain IRs) (23–25). Variable methyltransferase expression results in genome-wide methylation differences within a bacterial population, resulting in gene expression changes via epigenetic mechanisms; these systems are known as a phasevarions (<u>phase</u>-<u>vari</u>able regul<u>on</u>) (26, 27). In every case where a DNA methyltransferase has been shown to be phase-variable, the resulting phasevarion controls expression of multiple genes, with these genes often having roles in host colonisation and virulence, including antibiotic resistance, and many phasevarions regulate expression of putative vaccine candidates. Most phasevarions described to date are controlled by the ON-OFF switching of adenine specific (^m6^A) Type III DNA methyltransferases, encoded by *mod* genes. Phasevarions controlled by Type III Mod proteins have been described in a number of host-adapted bacterial pathogens such as non-typeable *Haemophilus influenzae* (27, 28), pathogenic *Neisseria* (26), *Helicobacter pylori* (29), and *Moraxella catarrhalis* (30). These Mod proteins all switch expression due to the variation in length of an SSR tract located in the open-reading frame of the encoding *mod* gene (Figure 1A). The DNA sequence recognised and methylated by Type III Mod proteins is determined by the target recognition domain (TRD). All currently characterised phase-variable *mod* genes can encode multiple different TRD regions, encoding multiple different Mod alleles in a species of bacteria . Varying the TRD results in a different DNA sequence being methylated, and consequently a different phasevarion being regulated. For example, there are currently 21 different *modA* alleles present in *H. influenzae* (28, 31) and the pathogenic *Neisseria* (26,31,32), 19 different *modH* alleles in *H. pylori*, (29, 33) and six *modM* alleles in *M. catarrhalis* (30).

**Figure 1.**
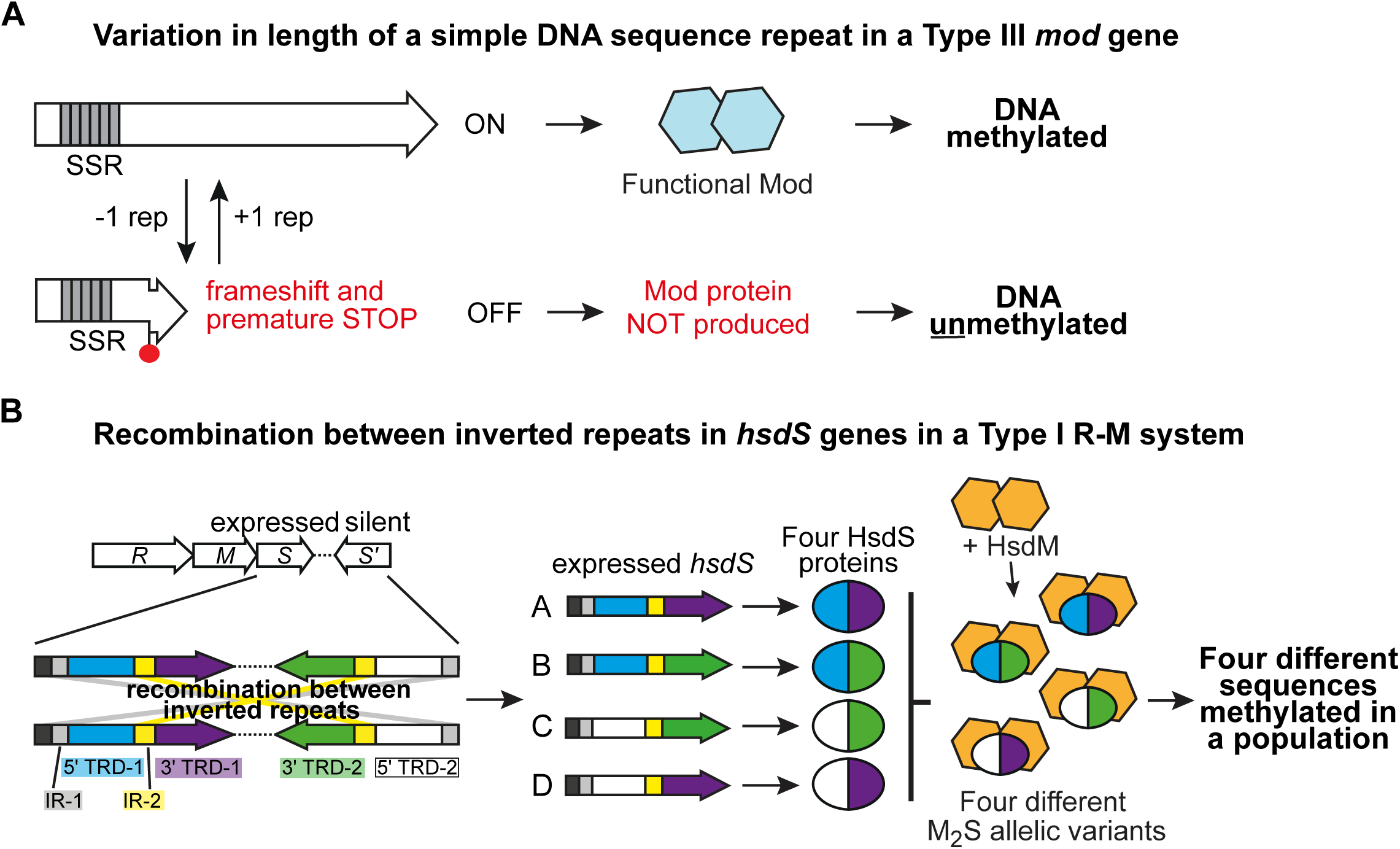
An illustration of the mechanisms of phase variation that occur in bacterial DNA methyltransferases. (A) Type III *mod* genes can contain variable length simple DNA sequence repeat (SSR) tracts in their open-reading frame. During DNA replication, polymerase slippage can occur when replicating these tracts, leading to loss or gain of repeat units. This variation in length results in the gene being in-frame downstream of the SSR tract, and expressed (ON), or a change in the reading frame leading to a frame-shift, a premature stop codon, and no expression of the encoded Mod protein (OFF). This means DNA is methylated (Mod ON) or not (Mod OFF); (B) Type I R-M systems that phase-vary often do so by shuffling between multiple variable *hsdS* specificity genes at the Type I R-M locus via recombination between inverted repeats (IRs) encoded in each of these multiple variable *hsdS* genes. This results in multiple different HsdS specificity proteins being expressed in a bacterial population, meaning multiple different DNA sequences are methylated in the bacterial population by the M _2_S trimer formed, dependent on the HsdS variant present in each individual bacterial cell of the population (four variants in the example in the illustration).

Type I DNA methyltransferases can also be phase-variably expressed. The first example of a phasevarion in a Gram-positive bacterial pathogen, *Streptococcus pneumoniae*, is controlled by a Type I DNA methyltransferase (34). The phasevarion present in *S. pneumoniae*, the SpnD39III system, switches between six different specificity subunits, encoded by *hsdS* genes, encoding HsdS proteins, which dictate the sequence restricted and methylated by the Type I system. The restriction and methyltransferase components of Type I systems are encoded by *hsdR* and *hsdM* genes, respectively (Figure 1B) (35). An active restriction enzyme is formed by an R_2_M2S pentamer, with a stand-alone methyltransferase formed by an M_2_S trimer. Each HsdS protein is made up of two individual TRDs, each of which contributes half to the overall specificity of the HsdS protein; therefore, changing a single TRD will result in a variation in the specificity of the resulting methyltransferase (Figure 1B). The SpnD39III system in *S. pneumoniae* contains multiple variable *hsdS* loci which contain inverted repeats, and a gene encoding a recombinase (*creX*). Homologous recombination between these multiple variable *hsdS* genes via these inverted repeats, and catalysed in part by the associated recombinase (36), results in the expression of six different methyltransferase variants within a population, which control six different phasevarions, resulting in six different phenotypic sub-variants within a pneumococcal population (34,37,38). A phase-variable Type I DNA methyltransferase has also been characterised in the related organism *Streptococcus suis* (39), with this system switching between four different HsdS variants. *S. suis* also encodes multiple phase-variable Type III methyltransferases which switch ON-OFF, and encoded by *modS* genes (40). Each distinct ModS allelic variant (ModS1-3) methylates a distinct DNA sequence, and controls a different phasevarion (40). *S. suis* therefore contains both phase-variable Type I and Type III DNA methyltransferases, although these systems show a discrete lineage distribution, with strains encoding either the phase-variable Type I system, or one of the three Type III ModS alleles (39). Our analysis of REBASE revealed the presence of methyltransferase loci with the hallmarks of phase variation in *A. pleuropneumoniae*: multiple *mod* genes that contained variable numbers of SSRs in their open-reading frame (23); and Type I systems encoding multiple variable, inverted *hsdS* loci containing inverted repeats, which would allow gene shuffling through homologous recombination (25). Therefore, we sought to characterise these methyltransferases in *A. pleuropneumoniae* and determine whether they controlled phasevarions. Defining these phasevarions allows us to identify the stably expressed antigenic repertoire of *A. pleuropneumoniae*, informing vaccine development against this major veterinary pathogen.

## MATERIALS AND METHODS

### Reagents

Genomic DNA was prepared using the Sigma GenElute bacterial genomic DNA kit according to the manufacturer’s instructions. Oligonucleotide primers were purchased from Integrated DNA Technologies (Singapore). PCR was carried out using GoTaq polymerase (Promega, Madison, USA) or KOD HotStart DNA polymerase (Merck, Darmstadt, Germany) according to the manufacturer’s instructions. Plasmid DNA was prepared using the Promega Wizard Mini Prep kit (Madison, USA). Restriction enzymes were purchased from New England Biolabs (NEB, Ipswich, MA, USA). DNA ligase, polynucleotide kinase (PNK) and Antarctic phosphatase used during cloning were all purchased from NEB (Ipswich, MA, USA). Purification of PCR products was carried out using the Qiagen PCR purification kit according to the manufacturer’s instructions.

### Bacteria and growth conditions

*A. pleuropneumoniae* isolates were grown on BBL™ blood agar base supplemented with thiamine HCl (0.0005%), NADH (0.0025%), oleic acid bovine albumin complex (5%) consisting of 4.75% bovine serum albumin in normal saline with heat inactivated horse serum (1 %), NaOH (5%) and oleic acid (0.06%) (termed BA/SN) (41) or brain heart infusion (BHI, Oxoid) agar containing β-nicotinamide adenine dinucleotide, NAD (0.01%) at 37°C for 24 h (42).

*Escherichia coli* strains (DH5α and BL21) were grown overnight in Luria-Bertani (LB) broth supplemented with ampicillin at 100 µg/ml or chloramphenicol at 20 µg/ml as required at 37 °C in a shaker at 200 rpm (39). For mutant selection, *A. pleuropneumoniae* strains were grown overnight on BHI-NAD plates with 1 µg/ml of chloramphenicol. The Type III methyltransferase genes were cloned into the pET15b vector. The prototype strains of *A. pleuropneumoniae* used in this study are listed in Table 1.

**Table 1.** Strains of *A. pleuropneumoniae* used in this study.

| Strains | Serovar | Phase-variable methyltransferase locus | accession number |
| --- | --- | --- | --- |
| AP76 | 7 | *modP1*, Type I | CP001091.1 |
| JL03 | 3 | *modP2* and *modQ*, Type I | CP000687.1 |
| S1536 | 2 | *modP3*, Type I | CP031875.1 |
| Femø | 6 | *modP4*, Type I | CP069796.1 |
| HS 143 | 15 | *modP1*, *modQ* | CP031859.1 |
| HS 3271 | 5 | *modP2* | No genome sequence available |

### Phylogenetic analysis

A total of 210 previously sequenced genomes, and all publicly available genomes from the RefSeq database in NCBI GenBank were used to determine the distribution of Type I and Type III R-M systems in *A. pleuropneumoniae* (Supplementary Table S1). We used the conserved *hsdR* and *hsdM* region of the Type I RM system from AP76 (GenBank locus tags APP7_1526-APP7_1531) (25), and prototype Type III *modP1-4 and modQ* genes from AP76 (GenBank locus tags APP7_0747), JL03 (GenBank locus tags APJL_0704) (Table 1). A tblasn search against a custom blast database of our collection of *A. pleuropneumoniae* was used to score the presence or absence of each system, where presence was determined by a >90% nucleotide identity over 90% of the length of the gene. To create a core genome alignment and phylogeny of our collection, we first annotated open reading frames using Prokka (*v1.12*) (43). Then we used the pangenome software Panaroo (*v1.2.7*) (44) with default parameters to create a core genome alignment. A neighbour joining tree was created from this alignment using the functions *dist.dna* and *nj* in the R package *ape* (45, 46) The distribution of the Type I and III systems were annotated on to the phylogeny using the interactive Tree of Life (iTOL) (47).

### Allelic diversity of Type III *mod* genes

An allele specific multiplex PCR was developed based on the TRD region of the prototype strains to identify different Type III methyltransferase genes and alleles present in un- sequenced isolates. A total of 250 field isolates of *A. pleuropneumoniae* submitted from 2000 to 2019 to the reference identification and serotyping service provided by the Microbiology Research Group (UQ, Brisbane, Australia) and serovar 1 to 15 *A. pleuropneumoniae* reference serovar strains were screened for Type III *mod* genes and their allelic variants (Supplementary Table S2) (48, 49). Multiplex PCR was carried out using GoTaq DNA polymerase (Promega) according to manufacturer’s instructions using the primers Mod-TRD- P1-F and Mod-TRD-P1-R, Mod-TRD-P2-F and Mod-TRD-P2-R, Mod-TRD-P3-F and Mod- TRD-P3-R, Mod-TRD-P4-F and Mod-TRD-P4-R, Mod-TRD-Q-F and Mod-TRD-Q-R, listed in Supplementary Table S3. Cycle conditions were as follows: initial denaturation at 95 °C for 2 min, followed by 30 cycles of denaturation at 95 °C for 30 s, annealing at 63 °C for 30 s, extension at 72 °C for 1 min, with a final extension at 72 °C for 10 min. Samples were checked on 1.5% (w/v) agarose gels buffered with 1x Tris-borate-EDTA.

### Fragment length analysis of Type III *mod* gene repeat tracts, and enrichment of ON-OFF strains

To quantify the ON-OFF ratio of each Type III *mod* gene, a *mod* specific primer pair was designed to amplify over the simple sequence repeat (SSR tract), using methodology as previously described (39). Briefly, a gene specific primer pair was designed for each of the two *mod* genes present (App-ModP-F-FAM and App-ModP-R or App-ModQ-F-Yakyel and App-ModQ-R, Supplementary Table S3) with the forward primer labelled with a fluorescent label to allow quantification using fragment length analysis (Life Technologies Australia Pty Ltd). Enrichment of strains expressing each Type III *mod* variant to generate populations of bacteria containing ∼90% ON or ∼90% OFF was carried out by single colony screening and enrichment (28). PCR was carried out using Go-Taq DNA polymerase (Promega) according to manufacturer’s instructions with following cycling conditions: initial denaturation at 95 °C for 2 min, followed by 30 cycles of denaturation at 95 °C for 30 s, annealing at 52 °C for 30 s, and extension at 72 °C for 30 s, with a final extension at 72 °C for 5 min. PCR fragments were sent to Australian Genome Research Facility (AGRF, Brisbane, Australia) and analysed with Peak Scanner Software v1.0 (Life Technologies Australia Pty Ltd).

### RNA extraction and cDNA synthesis

Triplicate cultures of *A. pleuropneumoniae* enriched ModP and ModQ ON/OFF variants were grown to mid-log (OD_600_ ∼ 0.4-0.7) in BHI broth supplemented with NAD (0.01%). RNA was prepared from 2 ml of bacterial culture using Trizol reagent (Thermo-Fisher) according to the manufacturer’s instructions. This RNA was used to synthesize cDNA using Protoscript II Reverse Transcriptase (NEB) and random hexamers (NEB) according to the manufacturer’s instructions. Reverse transcriptase reactions without Protoscript II was used as a negative control.

### Semi-quantitative RT-PCR

Semi-quantitative RT-PCR was carried out using GoTaq DNA polymerase (Promega) with 16S rRNA primers (16S F and 16S R) and one of four Type I allele - specific primer pairs (alleles A, B, C, D, E, F and G) (Supplementary Table S3) in a multiplex reaction using following cycling conditions: 95 °C for 2 min, followed by 30 cycles at 95 °C for 30 s, annealing at 54 °C for 20 s, extension at 72 °C for 45 s and final extension at 72 °C for 5 min (39). Samples were checked on 1.5% (w/v) agarose gels buffered with 1x Tris-borate-EDTA.

### Type I allele quantification

Allelic variants of each h*sdS* gene in *A. pleuropneumoniae* strain AP76 and JL03 were quantified using FAM labelled PCR coupled to a restriction digestion, based on methods as previously described (34, 39). Specifically, PCR amplification of the whole *hsdS* locus (3.7 kb) was carried out from extracted genomic DNA using GoTaq DNA polymerase (Promega) with primers App_T1_For_FAM and App_T1_Rev (Supplementary Table S3) according to manufacturer’s instructions. PCR cycle conditions were as follows: initial denaturation at 95 °C for 2 min, followed by 30 cycles of denaturation at 95 °C for 30 s, annealing at 52 °C for 30 s, and extension at 72 °C for 4 min, with a final extension at 72 °C for 5 min. The PCR product was then subjected to restriction digestion with BccI and NheI, with the digestions predicted to produce different sized fragments for each of the variant forms (Supplementary Figure S1). DNA fragments were sent to the Australian Genome Research Facility (AGRF, Brisbane, Australia) or the Griffith University DNA Sequencing Facility (GUDSF) and analysed with Peak Scanner Software v1.0 (Life Technologies Australia Pty Ltd). The area of the peak given by each labelled fragment, each corresponding to the prevalence of one of the variant forms, was quantified using Peak Scanner v1.0.

### Cloning and over-expression of Type III *mod* genes (*modP1, modP2* and *modQ*)

The *modP* (*modP1* and *modP2*) and *modQ* genes were PCR amplified using primers ModP- oE-F and ModP-oE-R, ModQ-oE-F and ModQ-oE-R (Supplementary Table S3) with KOD Hot- start DNA polymerase (Merck) according to manufacturer’s instructions. The resulting product was cloned into the NdeI and BamHI site of the pET15b vector and designated as pET15b::*modP1*, pET15b::*modP2*, and pET15b::*modQ*. After confirmation of the correct construct by colony PCR (primers: T7 forward and App-ModP-long for ModP and T7 forward and App-ModQ-long for Mod Q; Supplementary Table S3) and restriction digestion, proteins were over-expressed in *E. coli* BL21 at 37 °C with Isopropyl β-D-1-thiogalactopyranoside (IPTG) induction (0.5 mM) with 200rpm shaking. A no induction control was performed under identical conditions. The presence of over-expressed protein was confirmed using monoclonal anti-poly histidine alkaline phosphatase antibody (Sigma-Aldrich) at 1:20,000 dilution.

### Construction of modP1, modP2 and modQ mutants in A. pleuropneumoniae

All Mod knockout stains were constructed by replacing the TRD region with a chloramphenicol acetyltransferase gene (*cat*) (50), previously used to make insertional knockouts in *A. pleuropneumoniae* (51). The TRD region was first removed from the pET15b::*modP1* and pET15b::*modQ* by inverse PCR using the primers ModP_TRD_remove_F and ModP_TRD_remove_R; ModQ_TRD_remove_F and ModQ_TRD_remove_R, (Supplementary Table S3) and the amplified *cat* gene (Cm_USS_For and Cm_USS_Rev, Supplementary Table S3) was inserted by blunt end ligation. The constructs created were linearized and transformed into representative *A. pleuropneumoniae* strains via MIV transformation (52). Successful transformants were confirmed by PCR (App-ModP-F-FAM and CM-F-check; App-ModQ-F-Yakyel and CM-F-check, Supplementary Table S3) and sequencing.

### Construction of TRD-less Mod proteins

TRD-less Mod proteins were constructed via inverse PCR with primers ModP_TRD_remove_F and ModP_TRD_remove_R; ModQ_TRD_remove_F and ModQ_TRD_remove_R (Supplementary Table S3) using the vector pET15b::*modP1* and pET15b::*modQ* as a template. The PCR reaction was carried out using KOD Hot Start DNA polymerase (Merck) according to manufacturer’s instruction. The conserved 5 ′ and 3′ region was then fused and the TRD-less proteins over-expressed using *E. coli* BL21. Over- expression was carried out overnight at 37 °C (200 rpm) using IPTG (0.5 mM). After over- expression, proteins were purified using Talon resin (Takara Bio) by resuspending the cell pellet in lysis buffer (50 mM phosphate buffer and 300 mM NaCl) containing 0.1 mg/ml lysozyme and 0.5% SDS and multiple rounds of sonication. Proteins were eluted from the resin using gradient imidazole concentration (10 to 250 mM) in 1x binding buffer. After analysing fractions by SDS-PAGE and Western blot, pure fractions were concentrated and buffer exchanged using centrifugal concentrators (Amicon). TRD-less Mod proteins for both ModP and ModQ were then further purified by electroelution (100 mA for 30 min) into 1x MOPS buffer using a mini whole gel eluter (Biorad, Gladesville, NSW, Australia). Protein fractions were collected from each well and analysed by silver staining (Sigma) and Western blot using monoclonal anti poly histidine alkaline phosphatase antibody (1:20,000 dilution). Pure Mod TRD-less proteins (ModP and ModQ) were dialysed overnight at 4 °C in 1x PBS twice.

### Generation of Mod anti-sera

Anti ModP and ModQ sera (primary mouse antibody) were generated by immunizing five female BALB/c mice (6–8-week-old). A 50 µg aliquot of electroeluted purified proteins (TRD- less ModP and ModQ proteins) in alum were administered sub-cutaneously per mouse on day 0, 14, 28, 42. Terminal bleeds were collected on day 58. Serum was separated by centrifugation and stored at -20 °C in 50% glycerol. These antibodies (anti-ModP and anti- ModQ sera) were used as primary antibody. All animal work was approved by Griffith University Animal Ethics Committee (GLY/16/19/AEC).

### Western blotting

Whole cell lysates of each of the enriched Mod variants were prepared from mid-log (OD_600_ ∼0.4-0.7) cultures grown in BHI-NAD broth. Samples were normalized to OD_600_ of 2.0 and prepared for SDS-PAGE using Novex Bolt^TM^ LDS sample buffer and β-mercaptoethanol (final concentration 10% [v/v]). Samples were boiled for 20 min at 95 °C and centrifuged at 14,000x g for 20 min. Samples were run at 150 V for 1 hr on Novex Bolt™ Bis-Tris Plus precast Gels (4-12%) with 1x MOPS buffer (Life Technologies Australia Pty Ltd) and transferred to a nitrocellulose membrane (Bio-Rad, California, United States) at 15 V for 90 min. The membrane was then blocked overnight with 10% (w/v) skim milk in 1x Tris-buffered saline with Tween 20 (0.05%) (TBST) with shaking. The membrane was washed with 1x TBST 3-5 times for a total of 20 min, then incubated with primary mouse antibody raised against TRD-less ModP and ModQ proteins at 1:1,000 dilution for 1 hr at room temperature. The membrane was washed 3-5 times with 1x TBST for a total of 20 min and then incubated for an hour with 1:3,000 dilution of secondary antibody (goat anti-mouse alkaline phosphatase conjugate, Sigma). The membrane was washed again with 1x TBST 3-5 times for a total of 20 min and developed using 5-bromo-4-chloro-3-indolylphosphate (BCIP)/ nitro-blue tetrazolium (NBT) (Roche) according to the manufacturer’s instructions.

### Single-Molecule, Real-Time (SMRT) sequencing and methylome analysis

SMRT sequencing and methylome analysis was performed as described previously (53, 54) using genomic DNA isolated using the Sigma GenElute kit, or plasmid DNA isolated with the Qiagen miniprep kit. DNA was sheared (plasmid: ∼0.5–2kb; gDNA: 5–10 kb) using g-TUBEs (Covaris, Woburn, Massachusetts, USA) and sheared DNA was used to prepare SMRTbell template sequencing libraries. After end repairing, DNA was ligated to hairpin adapters and a combination of Exonuclease III (NEB; Ipswich, Massachusetts, USA) and Exonuclease VII (USB; Cleveland, USA) were used to degrade incompletely formed SMRTbell templates. Primer was annealed and samples were sequenced for long insert libraries on the PacBio Sequel system. SMRT sequencing and methylome analysis was carried out at SNPSaurus (University of Oregon, USA).

### Nanopore sequencing and methylome analysis

Genomic and plasmid DNA was isolated as above for SMRT sequencing. Nanopore sequencing libraries were prepared from 400 ng of DNA per sample with the Rapid DNA Barcoding kit (RBK004) according to manufacturer’s instructions. A 500 ng pool of 6 indexed plasmid libraries was sequenced on 1 PromethION R9.4.1 flow cells (FLO-PRO002) on a PromethION 24 instrument running MinKNOW 22.03.4. Reads were base called in super-accuracy mode (dna_r9.4.1_450bps_sup_prom) with demultiplexing, using Guppy 6.0.7 and default parameters. Read depth was downsampled to 400x per plasmid with ont_fast5_api v4.0.1 (https://github.com/nanoporetech/ont_fast5_api), first filtering for read quality with Filtong v0.2.1 (https://github.com/rrwick/Filtlong). DNA modification motif discovery and classification was performed with Nanodisco v1.0.3 (55), down-sampling read depth to 400x per plasmid, aligning reads to the pET15b vector containing the relevant *mod* insert. For genomic DNA libraries of *E. coli* and *A. pleuropneumoniae*, a 500 ng pool of 10 indexed samples was sequenced on 1 PromethION R9.4.1 flow cells (FLO-PRO002) on a PromethION 24 instrument running MinKNOW 22.08.6. Reads were basecalled in high-accuracy mode (dna_r9.4.1_450bps_hac_prom) with demultiplexing, using Guppy 6.1.5 and default parameters, and downsampled to a read depth of 400x per sample. DNA modification motif discovery and classification was performed with Nanodisco v1.0.3, aligning reads to the *E. coli* and *A. pleuropneumoniae* complete genomes. During the motif discovery step, a p-value threshold of -log_10_(p) > 20 was used for peak sequence selection. A threshold of -log_10_(p) > 100 was additionally used in analysis of *A. pleuropneumoniae* samples.

### SWATH MS proteomics

Overnight cultures of enriched ON/OFF variants (ModP1, ModP2 and ModQ) were grown to mid-log phase (OD_600_ ∼ 0.4–0.7) and cultures were normalized to OD_600_ of 1. Cells were harvested by centrifugation (5,500x g for 5 min) and resuspended in 300 µl of 6 M guanidinium chloride, 50 mM Tris-HCl (pH 8.0) and 10 mM dithiothreitol (DTT). Proteins were then alkylated by addition of 25 mM acrylamide and incubated for 60 min at 37 °C with 500 rpm. Proteins were precipitated by addition of methanol:acetone (1:1) and kept overnight at -20 °C. The protein was pelleted (18,000x g, 10 min) and resuspended in 50 µl of 50 mM Tris-HCl containing 1 µg trypsin (NEB). After overnight incubation at 37 °C, the trypsin digested peptides were cleaned with ZipTips (Millipore) and SWATH-MS analysis was performed as described previously (56). Briefly, tryptic peptides were analyzed by liquid chromatographyelectrospray ionization-tandem mass spectrometry (LC-ESI-MS/MS) using a Prominence nanoLC system (Shimadzu) and a TripleTOF 5600 mass spectrometer with a NanoSpray III interface (Sciex). Peptides were separated on a Vydac Everest reversed-phase C18 high-performance liquid chromatography (HPLC) column at a flow rate of 1ml/min. A gradient of 10% to 60% buffer B over 45 min, with buffer A (1% acetonitrile and 0.1% formic acid) and buffer B (80% acetonitrile and 0.1% formic acid) was used. A mass spectrometry (MS)-time of flight (TOF) scan was performed from an m/z range of 350 to 1,800 for 0.5 s, followed by information-dependent acquisition of MS/MS of the top 20 peptides from m/z 40 to 1,800 for 0.05 s per spectrum, with automated CE (capillary electrophoresis) selection. Identical LC conditions were used for SWATH-MS. SWATH-MS of triplicate biological replicates was performed with the same MS-TOF scan, followed by high-sensitivity information-independent acquisition with m/z isolation windows with 1 m/z window overlap each for 0.1 s across an m/z range of 400 to 1,250. Collision energy was automatically assigned by Analyst software (AB Sciex) based on m/z window ranges. Proteins were identified by searching against *A. pleuropneumoniae* strain AP76 and JL03 (NCBI accession no. CP001091.1 and CP000687.1, respectively) and common contaminants with standard settings using ProteinPilot 5.0.1 (AB Sciex). False-discovery-rate analysis was performed on all searches. ProteinPilot search results were used as ion libraries for SWATH analyses. The abundance of proteins was measured automatically using PeakView (AB Sciex) with standard settings. Comparison of protein relative abundance was performed based on protein intensities or ion intensities using a linear mixed-effects model with the MSstats package in R. Proteins with ≥2.0-fold changes in abundance and with adjusted P values greater than ≤0.05 were considered differentially expressed.

### Outer membrane protein (OMP) preparations

Enriched ON/OFF variants of *A. pleuropneumoniae* were grown to mid log phase (OD_600_ ∼ 0.4 - 0.7) in BHI broth containing 0.01% NAD. The cells were centrifuged at 3,000x g for 15 min and resuspended in 10 mM Tris-HCl (pH 8.0). OMPs were prepared as described previously (57). Briefly, cells were sonicated and centrifuged at 6,500x g for 15 min. The supernatant was transferred to a new tube and sarkosyl was added to a final concentration of 1% (v/v). The mixture was incubated for 30 min at room temperature and ultracentrifuged at 15,000x g for 90 min. The pellets were resuspended in 10 mM Tris-HCl (pH 8.0) containing sarkosyl (1% v/v final concentration) and incubated for 30 min at room temperature and, then ultracentrifuged at 15,000x g for 90 min. The supernatant was discarded and the OMP- enriched pellet was resuspended in 100 µl of 10 mM Tris-HCl (pH 8.0). The protein concentration was measured using a Pierce^TM^ BCA protein assay kit (Thermo Scientific). A 5 µg aliquot of each OMP preparation was run using the Novex Bis-Tris pre-cast gel system with 1x MOPS running buffer. Ammoniacal silver staining was carried out to visualize gross differences of protein expression due to ModP and ModQ phase variation (58). Differences in protein expression of selected proteins were quantified with ImageJ software by comparing the selected band intensity to a control band in the respective ON vs OFF lanes.

### Minimal inhibitory concentration

The minimum inhibitory concentration (MIC) was performed by a standardised broth microdilution method in Muller Hinton broth (MHB) with 2% yeast extract and NAD (0.1%) (59). The antimicrobials used were ampicillin, penicillin, florfenicol, ceftiofur, tulathromycin, tilmicosin and tiamulin. *A. pleuropneumoniae* ATCC 27090 strain was used as the quality control strain as recommended by CLSI guidelines (59). Briefly, overnight cultures of *A. pleuropneumoniae* enriched *mod* ON-OFF strains were grown to mid-log phase (OD_600_ ∼ 0.4 - 0.7). Cultures were then diluted to an OD_600_ of 0.2 and 50 μl of cultures were loaded into 96- well plates containing two-fold serially diluted antibiotics (ampicillin, penicillin, florfenicol from 5-0.1 µg/ml; tiamulin, tulathromycin, tilmicosin from 128-0.25 µg/ml; and enrofloxacin from 2-0.003 µg/ml). Plates were incubated for 24 hours at 37 °C under 5% CO_2_. The MIC (µg/ml) was determined as the lowest concentration of antibiotic to inhibit bacterial growth. Each assay was performed in biological triplicate.

### Growth curve and biofilm formation assays

A growth curve assay was performed by diluting overnight cultures of enriched ON/OFF variants (*modP1*, *modP2* and *modQ*) to OD_600_ of 0.2. The cultures were then inoculated in triplicate into a 96-well plate and incubated at 37 °C with shaking. Absorbance was measured (OD_600_) every 15 min using a BMG Labtech Fluostar Optima plate reader for 18 hr. Biofilm formation assays was performed as described previously (60). Enriched ON/OFF variants of *modP1*, *modP2* and *modQ* were grown overnight on a BHI-NAD agar plate (0.01% NAD) at 37 °C. The bacteria were harvested and suspended in fresh BHI-NAD broth and standardized to an OD_600_ of 0.2, and 100 μl of this suspension was inoculated in triplicate into a 96 -well plate. After overnight incubation at 37 °C, the broth was removed from each well and the well washed with distilled water. A 100 μl aliquot of crystal violet (0.1%) was then added into each well and incubated for 15 min at room temperature. The wells were washed with water three times and dried for 30 min at 37 °C. Biofilms were disrupted with 100 μL of 70% ethanol and absorbance was measured at OD_590_ using a BMG Labtech Fluostar Optima plate reader.

### Statistical analysis

GraphPad Prism 5.0 (GraphPad Software, La Jolla, California) was used to generate graphs and statistics. Error bars represent standard deviation from mean values. P values were considered as significant at <0.05 (*), <0.01 (**) and <0.001 (***), assessed using Student’s t test (unpaired, two tailed). P values of <0.05 were considered not statistically different (NSD).

## RESULTS

### Distribution of the phase-variable methyltransferases in *A. pleuropneumoniae*

Our analysis of REBASE (22) revealed the presence of multiple Type I and Type III methyltransferases in *A. pleuropneumoniae* genomes with the hallmarks of phase-variation (23, 25): duplicated variable *hsdS* genes containing IRs, or simple DNA sequence repeat (SSR) tracts. We observed the presence of multiple, duplicated *hsdS* genes in a Type I R-M system, with these duplicated variable *hsdS* genes containing inverted repeats (IRs). This Type I system also contained a gene annotated as a recombinase/integrase. The *hsdR* and *hsdM* genes encoded by this system were highly conserved (>95% nucleotide identity) in all 15 strains in REBASE identified to contain this system (25). The observed Type I system may therefore phase-vary much like the well characterised phase-variable Type I R-M system in *S. pneumoniae* strain D39 (34), the SpnD39III system, and subsequently described in multiple additional strains of *S. pneumoniae* (38, 61). The prototype *A. pleuropneumoniae* strains AP76 (NCBI accession no. CP001091.1, locus tags APP7_1526-APP7_1531) and JL03 (NCBI accession no. CP000687.1, locus tags APJL_1433- APJL_1438) both encoded this system.

All the Type III *mod* genes we identified contained a simple DNA sequence repeat tract within the open reading frame of each *mod* gene (23). All previously characterised *mod* genes containing SSRs have been shown to be phase-variable, and to control a phasevarion (26,28–30,40). Analysis of the sequences of the Type III *mod* genes we identified in *A. pleuropneumoniae* demonstrated the presence of two separate, unrelated phase-variable *mod* genes, which we named *modP* and *modQ*. Both *modP* and *modQ* genes contained a GCACA_(n)_ SSR tract in their 5′ region, downstream of the annotated ATG start codon. Further analysis of these sequences demonstrated that *modQ* was represented by a single variant. The *mod*Q TRD was the same in all strains of *A. pleuropneumoniae* in GenBank with a closed, annotated genome (28 strains in total) where a *modQ* gene is present (e.g. in strain JL03, NCBI accession no. CP000687, locus tag APJL_0819). Analysis of all *modP* sequences from these same 28 strains revealed the presence of four distinct TRD sequences, indicating at least four separate allelic variants of the *modP* gene present in the *A. pleuropneumoniae* population, similar to that seen with multiple other *mod* genes (26,28,39,62). We named these alleles *modP1* (e.g. in strain AP76, NCBI accession no. CP001091.1, locus tag APP7_0747), *modP2* (e.g. in strain JL03, NCBI accession no. CP000687.1; locus tag APJL_0704)*, modP3* (e.g. in strain S1536, NCBI accession no. CP031875.1, locus tag D1095_03465), and *modP4* (e.g. in strain Femø, NCBI accession no. CP069796.1, locus tag D1099_04110).

Using conserved sequences of the identified Type I R-M system (the *hsdR* and *hsdM* regions), and each individual phase-variable Type III *mod* gene, we carried out a detailed sequence analysis using a diverse collection of 182 *A. pleuropneumoniae* isolates, and the 28 strains with a publicly available fully annotated whole genome to determine the distribution of each of these methyltransferases in *A. pleuropneumoniae* (Figure 2) (63). All 210 strains used for phylogenetic analysis are listed in Supplementary Table S1. This analysis demonstrated that all strains contained at least one of the Type III *mod* genes containing an SSR tract (19). Of these 210 strains, the *modP1* allele was present in 150 strains followed by *modQ* in 61 strains, *modP3* in 20 strains, *modP2* and *modP4* in 17 strains. Our analysis demonstrated that approximately 26% (n = 56) of strains harboured both *modP* and *modQ* genes. This analysis also revealed the distribution of Type III *mod* genes is serovar specific. For example, 100% of serovar 8 (n = 104) and serovar 7 (n = 21) encoded the *modP1* gene whilst the *modP3* gene was only present in serovar 2 (n = 20). All serovar 5 isolates (n = 6) were found to harbour the *modP2* gene. For serovar 12, six isolates encoded the *modP2* allele and one isolate encoded the *modP4* allele, which was also present in all serovar 6 isolates (n = 10). The *modQ* gene appears to be absent in all isolates of serovars 4, 7, 8, 9, 17, and K2:07, as well as 3/21 serovar 2 isolates, whereas it is present in all other serovar isolates (Figure 2, Supplementary Table S1). In addition, 195 strains (93%) encoded the Type I locus encoding duplicate, variable *hsdS* loci containing inverted repeats (64). Of these 195 strains, 78% (n = 152) of strains encoded both the Type I system and one of the Type III *mod* genes containing SSRs; 22% (n = 43) of strains had both the phase-variable Type I system and two Type III *mod* genes containing SSRs. This analysis indicated that individual strains can contain both phase-variable Type I and Type III methyltransferases, a phenomenon never before observed in individual bacterial strains.

**Figure 2.**
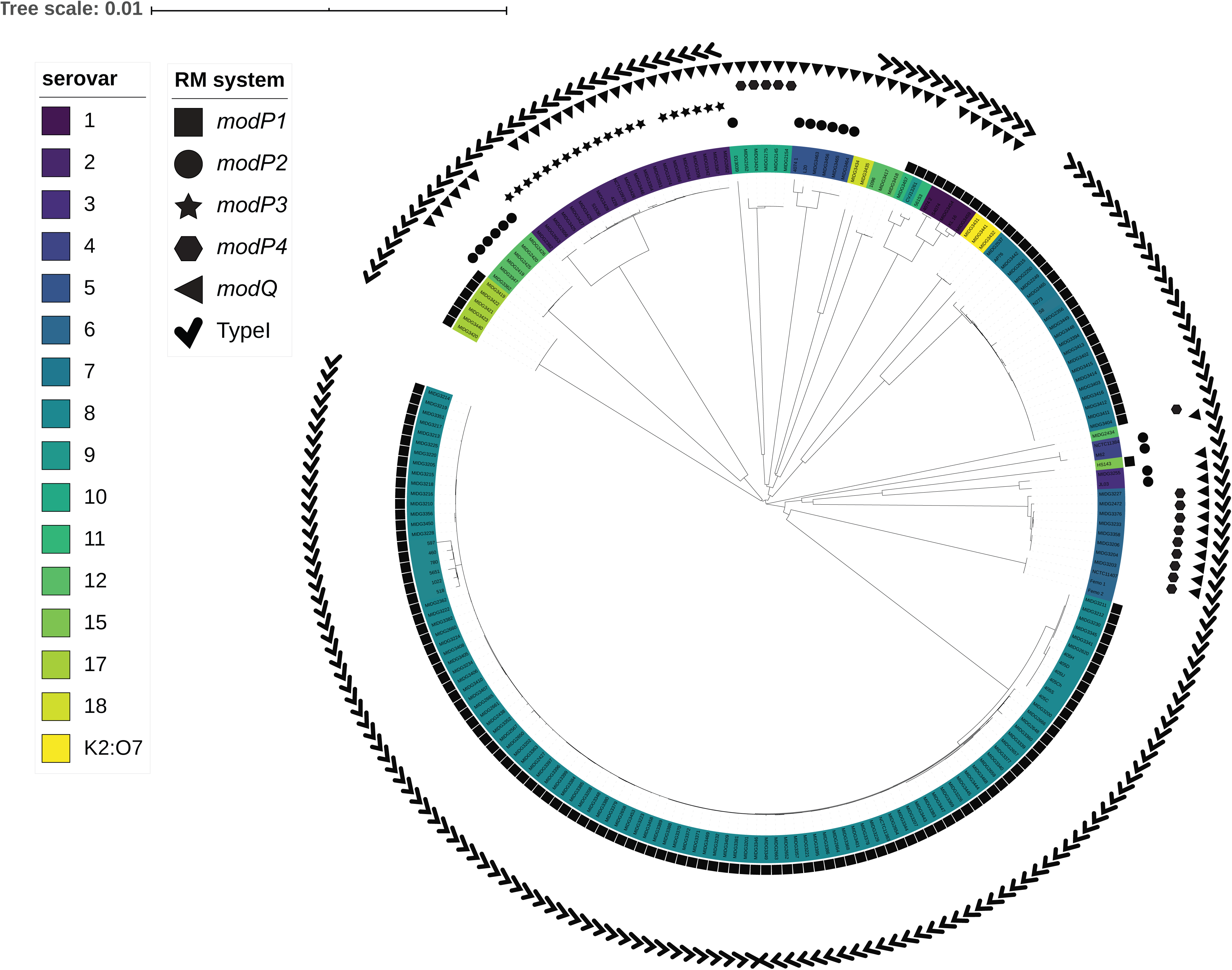
The distribution of phase-variable Type I and Type III methyltransferases in *A. pleuropneumoniae.* A neighbour joining tree of *A. pleuropneumoniae* showing the genetic distance between isolates based on all sites in a core genome. Serovars are represented by coloured clades and the presence of phase variable Type I and Type III methyltransferases are indicated by different shapes. The scale bar indicates the genetic distance (number of substitutions per site).

We also investigated the distribution of these newly identified Type III *mod* genes in a second collection of *A. pleuropneumoniae* strains, comprising 250 field isolates and 15 reference strains that represented one of each of the 15 serovars of *A. pleuropneumoniae* (65) (Supplementary Table S2). These field isolates comprise the Australian national reference collection for *A. pleuropneumoniae* (48, 49). These field isolates have not been whole genome sequenced, and represented isolates from diverse geographical locations around Australia and across the globe. To determine the Type III genes/alleles present in this collection, we designed primer pairs specific for the TRDs of each *modP* allele, and for *modQ*, in order to screen via PCR. This analysis showed that the most prevalent serovars in Australia (serovar 1, 5, 7, 12 and serovar 15) (66) harboured the same allelic variant(s) of these newly identified *mod* genes as in our first collection. For example, 100% of serovar 1 and serovar 15 isolates encoded the *modP1* allele and the *modQ* gene. The *modP1* allele and *modQ* gene were also present in *A. pleuropnuemoniae* reference strains of serovar 9 and 11. The *modP2* allele and the *modQ* gene were found in all serovar 5 isolates (100%). For serovar 7, almost all (96%, n = 67) isolates were found to contain only the *modP1* allele, with the exception of a single serovar 7 reference strain (WF83) and 2 Australian field isolates that contained the *modP1* allele and *modQ* gene. The *modP3* allele was present in serovar 2 (n = 2), 12 (n = 4), serovar 14 reference strain (n = 1) and two non-typeable strains. No strain in our culture collection encoded the *modP4* allele except our single serovar 6 reference strain, Femø. Analysis of the field isolates revealed that 97% isolates encoded the *modP* gene (n = 256), with the *modP1* allele being the most prevalent, present in ∼78% (n = 206) of all isolates. The *modP2* allele was found at a lower frequency, comprising only 15% of isolates (n = 39) and only 3.8% (n = 10) isolates encoded the *modP3* allele. In addition, 70% of *A. pleuropneumoniae* isolates in this collection (n = 185) contained the *modQ* gene. Overall, approximately 65% (n =173) of isolates contained at least one of the four *modP* alleles and the *modQ* gene. Due to high prevalence of *modP1*, *modP2* and *modQ* alleles in our Australian culture collections, we selected these three alleles for further analysis in this study.

### The Type I system observed in *A. pleuropneumoniae* is phase-variable and encodes multiple different TRDs resulting in different HsdS proteins in different strains

In order to determine the extent of variable HsdS proteins that are encoded by the almost ubiquitous phase-variable Type I R-M system we have observed in *A. pleuropneumoniae* (Figure 2; illustrated in Figure 3A), we analysed the sequences of all TRD regions available in REBASE (15 strains). Analysis of these TRD sequences revealed that there are four unique 5*′*-TRD sequences, and four unique 3*′*-TRD sequences present in these strains (Figure 3B; full sequences and analysis in Supplementary Data 1), meaning a potential 16 unique HsdS proteins were possible dependent on the encoded TRDs present (four possible 5*′*-TRD x four possible 3*′*-TRD). This is unlike other phase-variable Type I systems, such as the SpnD39III system in *S. pneumoniae* (34,37,38,61), and the phase-variable Type I system in *S. suis,* (39) where all strains encode the same TRDs, and therefore shuffle between the same complement of expressed HsdS proteins. Our prototype strains of AP76 and JL03 encoded a subset of these sixteen potential TRDs: AP76 encoded a combination of 5*′*-TRD-1 and 3*′*- TRD-1; 5*′*-TRD-1 and 3*′*-TRD-2; 5*′*-TRD-2 and 3*′*-TRD-2; 5*′*-TRD-2 and 3*′*-TRD-1, which we have named alleles A-D, respectively (Figure 3C). In strain JL03, we found *hsdS* allele D (5*′*- TRD-2 and 3*′*-TRD-1), along with three distinct *hsdS* alleles, encoded by the combinations 5*′*- TRD-2 and 3*′*-TRD-3; 5*′*-TRD-3 and 3*′*-TRD-3; and 5*′*-TRD-3 and 3*′*-TRD-1), which we have named alleles E-G, respectively (Figure 3D). In order to demonstrate that these *hsdS* alleles are phase-variable, we performed a semi-quantitative RT-PCR to demonstrate that all four potential *hsdS* allelic variants were expressed in a population of *A. pleuropneumoniae*, as carried out previously for a phase-variable Type I system in *S. suis* (39). This analysis demonstrated that all four *hsdS* alleles that we predicted to be present (alleles A-D in strain AP76, and alleles D-G in strain JL03) were expressed in a population of the respective strains (Figure 3C and 3D). This analysis provides excellent evidence that phase variation of this Type I system is occurring in both strains by shuffling of variable TRDs between the expressed and silent *hsdS* loci present. We also designed a FAM labelled PCR coupled to fragment length analysis to quantify the proportion of bacteria expressing each allele within a population of *A. pleuropneumoniae* (Supplementary Figure S1). Using these same prototype strains AP76 and JL03, we showed that the majority of the population in strain AP76 expressed *hsdS* allele A (5*′*-TRD-1 and 3*′*-TRD-1), and in strain JL03 *hsdS* allele E (5*′*-TRD-2 and 3*′*-TRD-3), but all other minor variants were also detectable using this methodology (Supplementary Figure S1D). Along with our semi-quantitative RT-PCR, this conclusively demonstrates phase variation of the Type I system in *A. pleuropneumoniae*.

**Figure 3.**
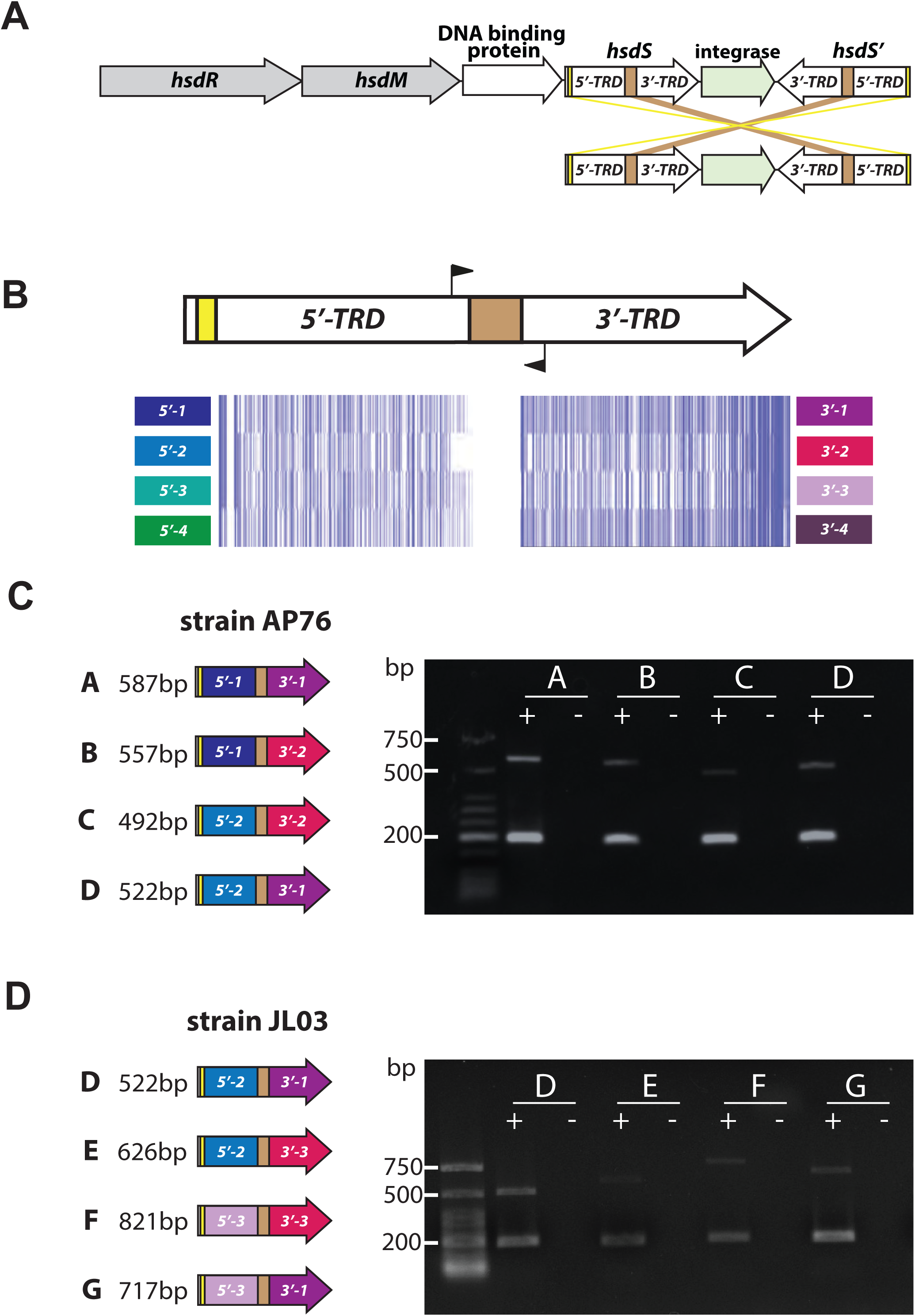
Demonstration of the phase-variable Type I *hsdS* locus. (A) Illustration of the Type I R-M system containing duplicate, variable *hsdS* specificity loci that contain IRs. Gene shuffling between variable 5’ and 3’ TRDs produces four unique *hsdS* genes at the *hsdS* expressed locus immediately downstream of the *hsdM* gene, results in the expression of four unique allelic variants of the HsdS protein; (B) Alignments of the four 5’ and four 3’ TRDs we present in 15 *A. pleuropneumoniae* strains available in REBASE (see full sequences and analysis in Supplementary Data 1). Alignments were carried out using Muscle and visualized in JalView overview feature; (C) and (D) Semi-quantitative RT-PCR was carried out to demonstrate the presence of cDNA and thus mRNA for each of the four encoded *hsdS* alleles, indicating that all four alleles are expressed in the population of AP76 and JL03. Expected sizes are: allele A = 587 bp, allele B = 557 bp, allele C = 492 bp, allele D = 522 bp, allele E = 626 bp, allele F = 821 bp, allele G = 717 bp.

### ModP is the first characterised phase-variable cytosine specific Type III DNA methyltransferase

As described above, we determined the presence of four unique *modP* alleles in *A. pleuropneumoniae*. Each allelic variant shows high sequence variability in the central TRD encoding region (illustrated in Figure 4A), with <25% nucleotide identity between each *modP* allele (Figure 4B), consistent with that seen for other phase-variable *mod* genes (39). The 5′ and 3′ regions of these *modP* alleles share >95% nucleotide sequence identity.

**Figure 4.**
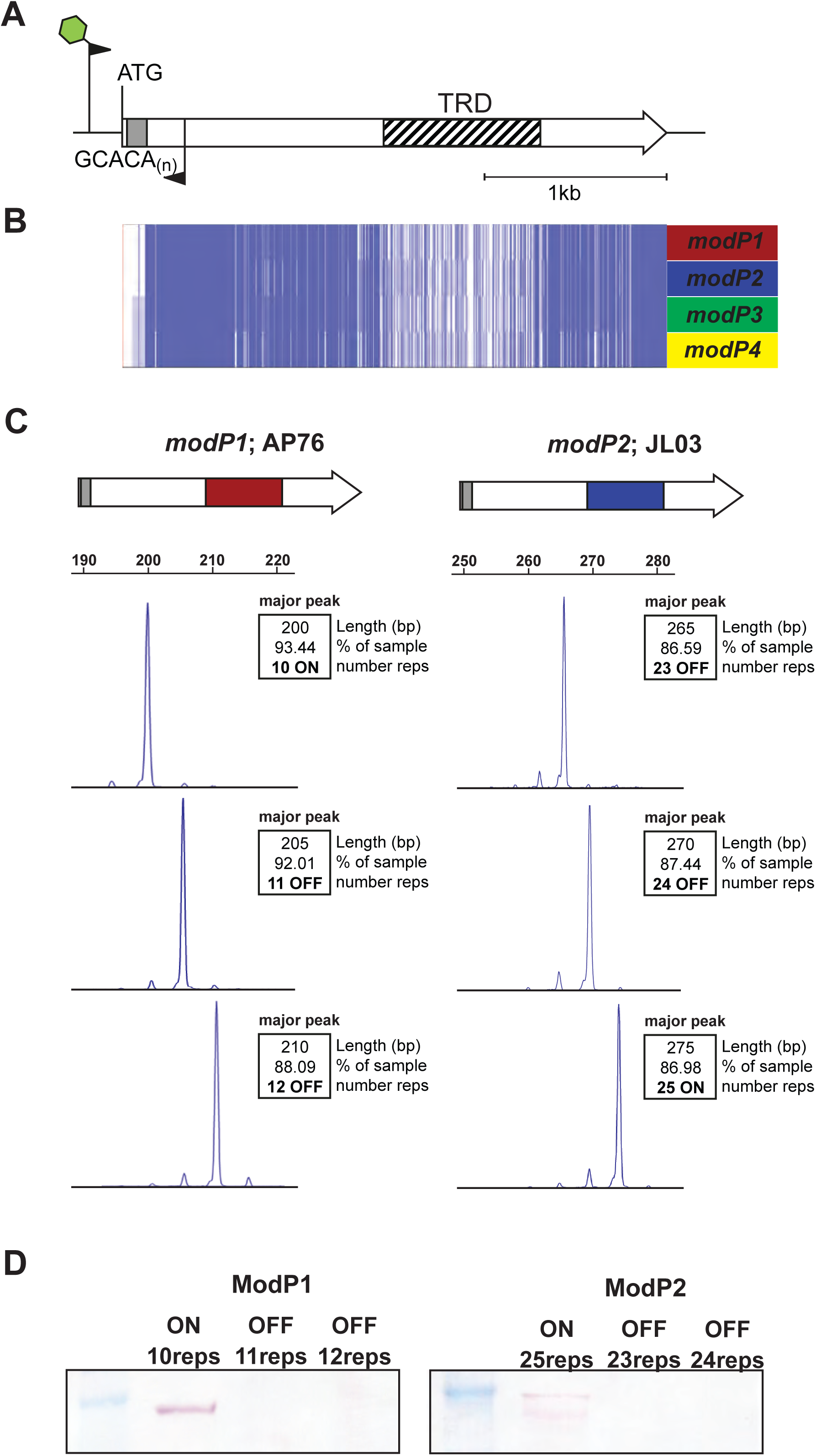
Phase-variable expression of *modP*. (A) Illustration of the Type III *modP* gene present in *A. pleuropneumoniae*. The *modP* gene contains a variable number of GCACA_(n)_ repeats immediately downstream of the ATG start codon, and a highly variable target recognition domain (TRD) that dictates DNA sequence specificity of the encoded protein, represented by the hatched box. The green star on the forward primer depicts a 6- Carboxyfluoresceine (FAM) fluorescent label which allows analysis of PCR products using GeneScan technology (Applied Biosystems International). (B) Full-length DNA sequences of *modP1, modP2, modP3* and *modP4* were aligned using Muscle and visualized in JalView overview feature. Each blue line represents one nucleotide; (C) Fragment length analysis PCR over the GCACA_(n)_ repeat tracts from enriched modP1 and modP2 strains with three consecutive SSR tract lengths in strains AP76 (*modP1*) and JL03 (*modP2*); (D) Western blot analysis using anti-ModP sera confirmed phase-variable expression of ModP, which was only present in a population of *A. pleuropneumoniae* enriched for 10 repeats (ModP1) or 25 repeats (ModP2).

To examine if these *modP* genes were phase-variably expressed, we performed single colony isolation and enrichment to isolate individual variants of prototype strains encoding the two most common alleles, *modP1* and *modP2*. This would allow enrichment for a triplet set of isogenic strains that were ∼90% pure for a single GCACA_(n)_ SSR tract length in the respective *modP* gene. We performed a FAM-labelled PCR coupled to fragment length analysis over the GCACA_(n)_ SSR tract to isolate these variants, each representing one of the three reading frames possible (Figure 4C). We used strain AP76 as our prototype strain for *modP1* and strain JL03 for *modP2*. This resulted in a triplet of *modP1* strains in AP76 where the GCACA_(n)_ SSR tract was 10, 11, or 12 repeats and for *modP2* in JL03 with 23, 24 or 25 repeats (Figure 4C). Using whole cell lysates of these enriched triplet sets of each prototype strain we carried out Western blotting with antisera raised against the conserved region of ModP. This demonstrated that each ModP variant is only expressed when the GCACA_(n)_ SSR tract maintains the open reading frame (ON), and not expressed when the SSR tract length results in a frameshift and premature stop codon (OFF) (Figure 4D, full blot in Supplementary Figure S2). ModP1 in strain AP76 was only expressed when there were 10 GCACA repeats present, and ModP2 in strain JL03 when there were 25 GCACA repeats present.

Using long read sequencing from both PacBio SMRT sequencing and Oxford Nanopore technology, we performed methylome analysis of both ModP1 and ModP2. We did this by sequencing both plasmid vectors and genomic DNA isolated from *E. coli* BL21 strains over- expressing each cloned methyltransferase, and genomic DNA isolated from each of our triplet sets of prototype *A. pleuropneumoniae* strains encoding either the *modP1* gene (strain AP76) or the *modP2* gene (strain JL03) and *modP1* and *modP2* knockout *A. pleuropneumoniae* mutants. This analysis demonstrated that both ModP1 and ModP2 are cytosine specific Type III DNA methyltransferases, each recognising and methylating a different DNA target sequence. ModP1 methylates a cytosine in the motif CAA**^(m4)^C**T, and ModP2 methylates a cytosine in the motif GWC(**^m4^)C**T. Cytosine specific Type III DNA methyltransferases have been previously identified (67), but the ModP methyltransferase is the first identified phase-variable cytosine-specific Type III DNA methyltransferase (Table 2; all methylome data is presented in Supplementary Data 2; examples of PacBio and Nanopore analysis for each type of DNA analysed and motifs detected is presented in Supplementary Figures S3-S6).

**Table 2.** Summary of methyltransferase specificities and methodology used to determine for ModP1, ModP2, and ModQ. Y = detected; nd = not detected; nc = method not carried out

|  | ModP1 | ModP2 | ModQ |
| --- | --- | --- | --- |
|  | CAA<sup>(m4)</sup>CT | GWC<sup>(m4)</sup>CT | AG<sup>(m6)</sup>ATG |
|  | Motif detected by: |  |  |
| SMRT sequencing *A. pleuropneumonia* gDNA | Y | nc | Y |
| Nanopore sequencing *A. pleuropneumonia* gDNA | nd | nd | Y |
| Nanopore sequencing *E. coli* gDNA | nd | Y | Y |
| SMRT sequencing *E. coli* plasmid DNA | nc | Y | Y |
| Nanopore sequencing *E. coli* plasmid DNA | nd | Y | Y |

### ModP1 and ModP2 regulate different phasevarions

Following this novel finding, we wanted to determine if ModP1 and ModP2 phase-variable expression resulted in the regulation of different phasevarions, i.e., did cytosine methylation at different motifs result in different sets of proteins being regulated commensurate with the observed phase variation? To quantify the extent of differential protein expression mediated by ModP1 and ModP2 phase variation, we carried out quantitative SWATH proteomics, used previously to characterise expression differences mediated by phase-variable DNA methyltransferases (40). The expression profiles of the ModP1 ON-OFF pair (10 repeats ON and 11 repeats OFF) and ModP2 ON-OFF pair (25 repeats ON and 24 repeats OFF) were assessed by SWATH-MS proteomics. Both strain pairs showed unique differences in protein expression dependent on which ModP protein was expressed, i.e., ModP1 and ModP2 control different phasevarions. Our SWATH-MS covered 26% of total proteins in strain AP76 (ModP1) and 32% of total proteins in strain JL03 (ModP2). In addition, gross protein expression differences were observed using outer membrane protein (OMP) fractions prepared from our enriched *modP1* and *modP2* ON-OFF pairs (Supplementary Figure S7).

In strain AP76 encoding ModP1, a total of 42 proteins had an expression ratio of ≥2.0 fold different between the ModP1 ON and ModP1 OFF isogenic strains. Twenty-seven proteins were upregulated in ModP1 ON, and fifteen proteins were downregulated in ModP1 ON (Table 3). Of the proteins showing increased expression in our ModP1 variant, many are involved in pathobiology, including the major RTX toxin, ApxIVA, a heat shock-like protein 33 HslO, and outer membrane protein W. Several proteins involved in energy metabolism were also upregulated in ModP1 ON including succinyl-CoA synthetase beta chain and mannose permease IID. In addition, several regulatory proteins were upregulated in ModP1 ON, such as leucine-responsive regulatory protein (Lrp), and a putative HTH-type transcriptional. Several biosynthetic proteins showed an increase in expression in ModP1 ON, including those involved in amino acid biosynthesis (Ketol-acid reductoisomerase, D-3-phosphoglycerate dehydrogenase, aspartate-semialdehyde dehydrogenase), heme biosynthesis (coproporphyrinogen III oxidase) and lipid A biosynthesis (acylUDP-N- acetylglucosamine O-acyltransferase). In the ModP1 OFF strain, many ribosomal proteins, outer membrane protein precursor PalA, lipoprotein and the fatty acid biosynthesis protein (Malonyl CoA-acyl carrier protein transacylase (MCT)) exhibited increased expression compared to ModP1 ON (Table 3).

**Table 3.**
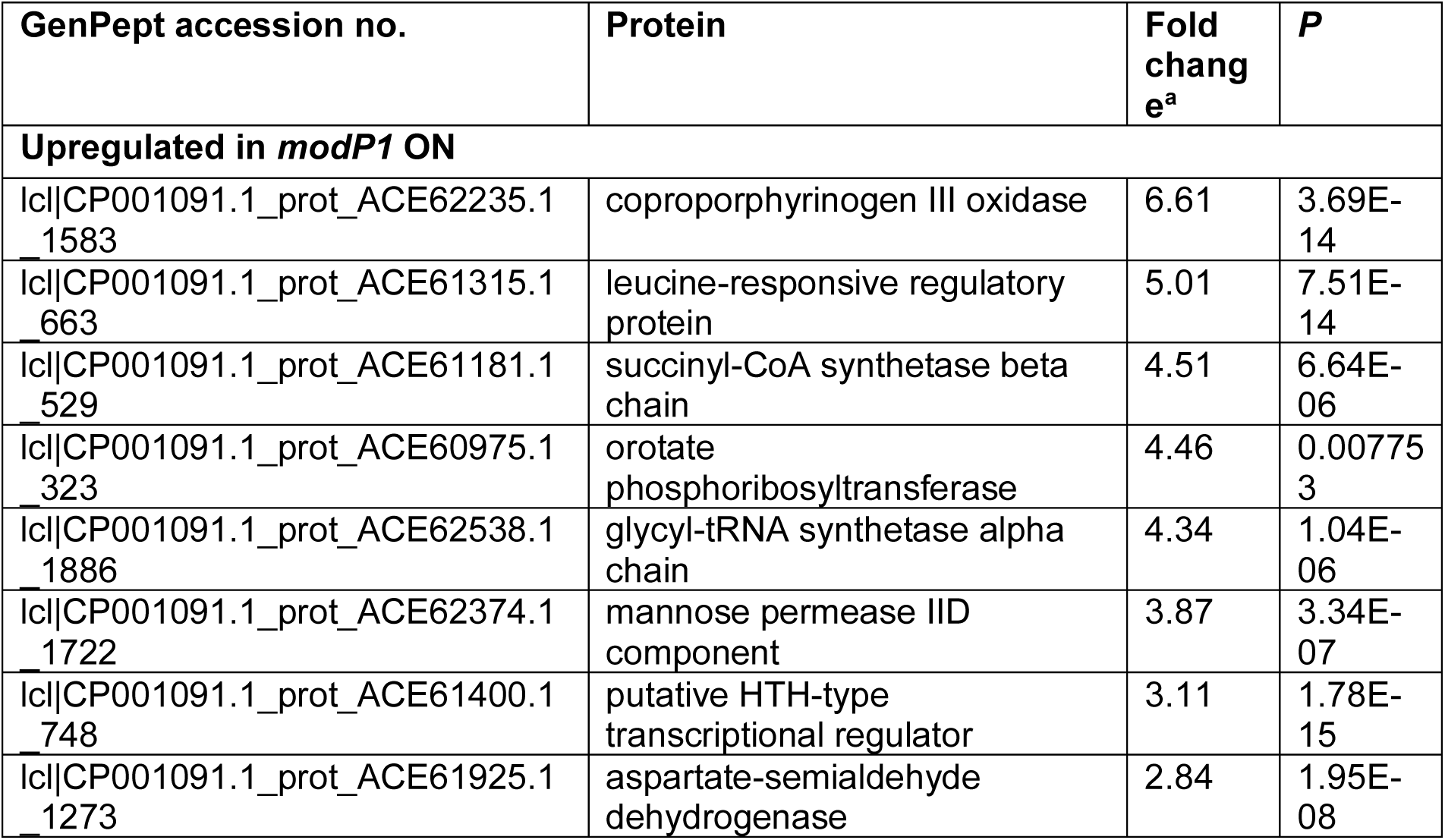
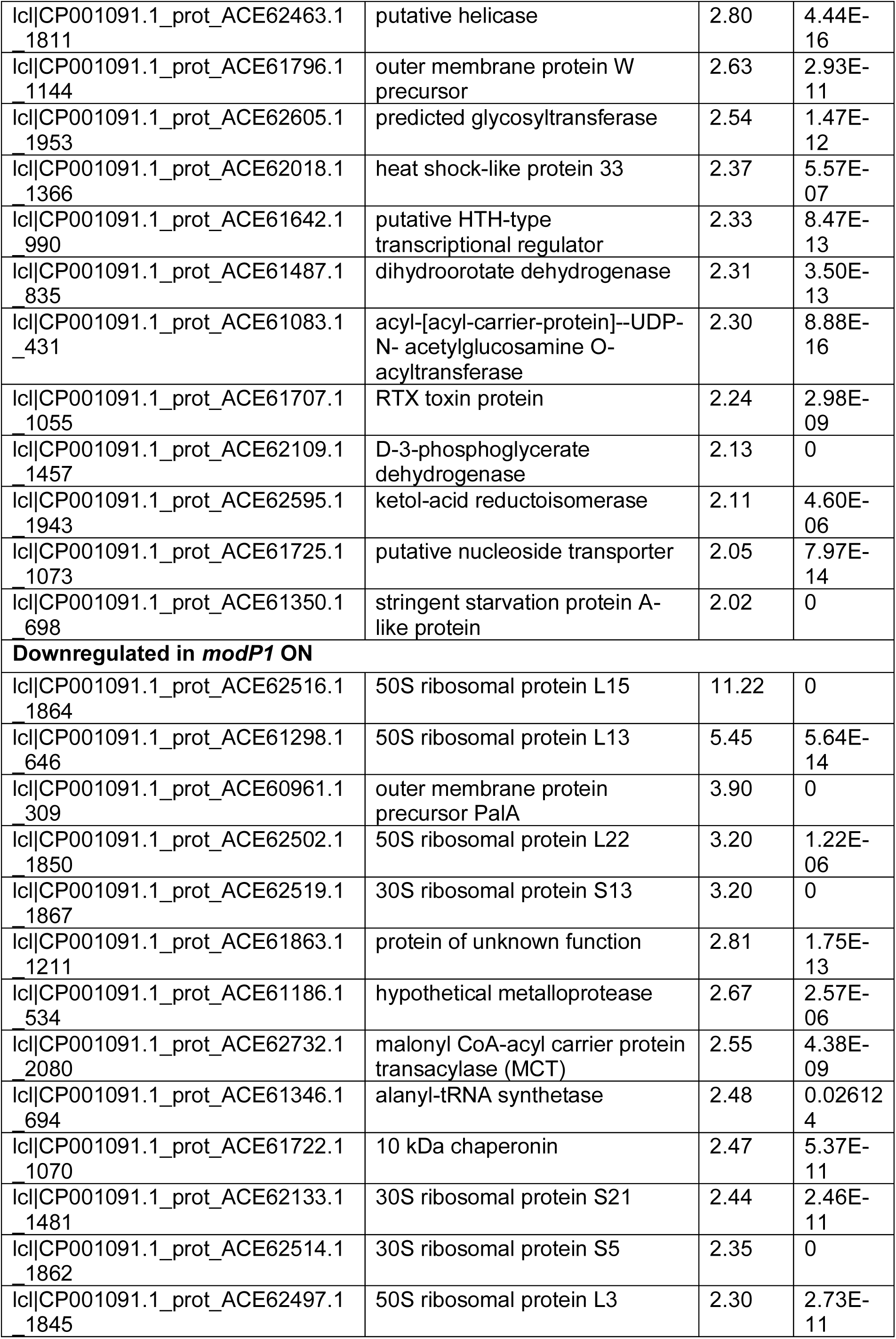
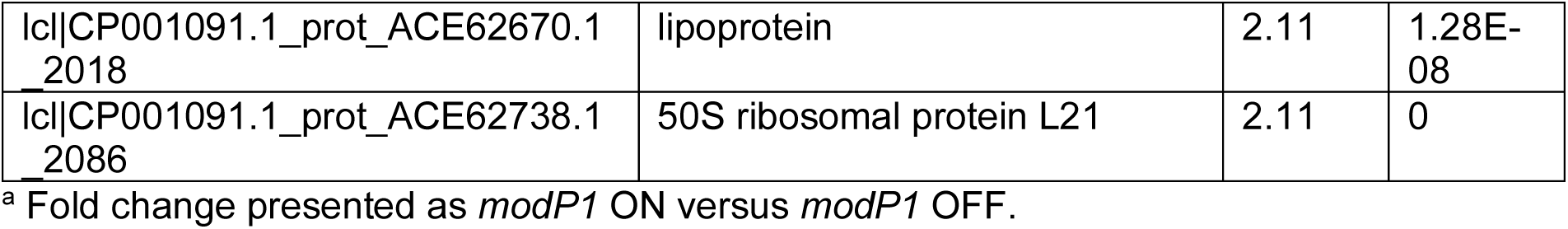
Differentially regulated proteins (≥2.0 fold) in the ModP1 phasevarion.

| GenPept accession no. | Protein | Fold change<sup>a</sup> | P |
| --- | --- | --- | --- |
| **Upregulated in *modP1* ON** |  |  |  |
| Ic\|CP001091.1_prot_ACE62235.1_1583 | coproporphyrinogen III oxidase | 6.61 | 3.69E-14 |
| Ic\|CP001091.1_prot_ACE61315.1_663 | leucine-responsive regulatory protein | 5.01 | 7.51E-14 |
| Ic\|CP001091.1_prot_ACE61181.1_529 | succinyl-CoA synthetase beta chain | 4.51 | 6.64E-06 |
| Ic\|CP001091.1_prot_ACE60975.1_323 | orotate phosphoribosyltransferase | 4.46 | 0.007753 |
| Ic\|CP001091.1_prot_ACE62538.1_1886 | glycyl-tRNA synthetase alpha chain | 4.34 | 1.04E-06 |
| Ic\|CP001091.1_prot_ACE62374.1_1722 | mannose permease IID component | 3.87 | 3.34E-07 |
| Ic\|CP001091.1_prot_ACE61400.1_748 | putative HTH-type transcriptional regulator | 3.11 | 1.78E-15 |
| Ic\|CP001091.1_prot_ACE61925.1_1273 | aspartate-semialdehyde dehydrogenase | 2.84 | 1.95E-08 |
| Icl\|CP001091.1_prot_ACE62463.1_1811 | putative helicase | 2.80 | 4.44E-16 |
| Icl\|CP001091.1_prot_ACE61796.1_1144 | outer membrane protein W precursor | 2.63 | 2.93E-11 |
| Icl\|CP001091.1_prot_ACE62605.1_1953 | predicted glycosyltransferase | 2.54 | 1.47E-12 |
| Icl\|CP001091.1_prot_ACE62018.1_1366 | heat shock-like protein 33 | 2.37 | 5.57E-07 |
| Icl\|CP001091.1_prot_ACE61642.1_990 | putative HTH-type transcriptional regulator | 2.33 | 8.47E-13 |
| Icl\|CP001091.1_prot_ACE61487.1_835 | dihydroorotate dehydrogenase | 2.31 | 3.50E-13 |
| Icl\|CP001091.1_prot_ACE61083.1_431 | acyl-[acyl-carrier-protein]--UDP-N- acetylglucosamine O-acyltransferase | 2.30 | 8.88E-16 |
| Icl\|CP001091.1_prot_ACE61707.1_1055 | RTX toxin protein | 2.24 | 2.98E-09 |
| Icl\|CP001091.1_prot_ACE62109.1_1457 | D-3-phosphoglycerate dehydrogenase | 2.13 | 0 |
| Icl\|CP001091.1_prot_ACE62595.1_1943 | ketol-acid reductoisomerase | 2.11 | 4.60E-06 |
| Icl\|CP001091.1_prot_ACE61725.1_1073 | putative nucleoside transporter | 2.05 | 7.97E-14 |
| Icl\|CP001091.1_prot_ACE61350.1_698 | stringent starvation protein A-like protein | 2.02 | 0 |
| **Downregulated in *modP1* ON** |  |  |  |
| Icl\|CP001091.1_prot_ACE62516.1_1864 | 50S ribosomal protein L15 | 11.22 | 0 |
| Icl\|CP001091.1_prot_ACE61298.1_646 | 50S ribosomal protein L13 | 5.45 | 5.64E-14 |
| Icl\|CP001091.1_prot_ACE60961.1_309 | outer membrane protein precursor PalA | 3.90 | 0 |
| Icl\|CP001091.1_prot_ACE62502.1_1850 | 50S ribosomal protein L22 | 3.20 | 1.22E-06 |
| Icl\|CP001091.1_prot_ACE62519.1_1867 | 30S ribosomal protein S13 | 3.20 | 0 |
| Icl\|CP001091.1_prot_ACE61863.1_1211 | protein of unknown function | 2.81 | 1.75E-13 |
| Icl\|CP001091.1_prot_ACE61186.1_534 | hypothetical metalloprotease | 2.67 | 2.57E-06 |
| Icl\|CP001091.1_prot_ACE62732.1_2080 | malonyl CoA-acyl carrier protein transacylase (MCT) | 2.55 | 4.38E-09 |
| Icl\|CP001091.1_prot_ACE61346.1_694 | alanyl-tRNA synthetase | 2.48 | 0.026124 |
| Icl\|CP001091.1_prot_ACE61722.1_1070 | 10 kDa chaperonin | 2.47 | 5.37E-11 |
| Icl\|CP001091.1_prot_ACE62133.1_1481 | 30S ribosomal protein S21 | 2.44 | 2.46E-11 |
| Icl\|CP001091.1_prot_ACE62514.1_1862 | 30S ribosomal protein S5 | 2.35 | 0 |
| Icl\|CP001091.1_prot_ACE62497.1_1845 | 50S ribosomal protein L3 | 2.30 | 2.73E-11 |
| lcl\|CP001091.1_prot_ACE62670.1<br>2018 | lipoprotein | 2.11 | 1.28E-08 |
| lcl\|CP001091.1_prot_ACE62738.1<br>2086 | 50S ribosomal protein L21 | 2.11 | 0 |

<sup>a</sup> Fold change presented as *modP1* ON versus *modP1* OFF.

In strain JL03, phase variation of ModP2 resulted in varied expression of a different set of proteins, with 35 proteins showing a ≥2.0 fold expression ratio between ModP2 ON and ModP2 OFF, of which 9 proteins were upregulated in ModP2 ON, and 26 proteins downregulated in ModP2 ON (Table 4). Our ModP2 ON strain showing increased expression of proteins involved in central metabolism, including an acyl carrier protein, a biotin carboxyl carrier protein of acetyl-CoA carboxylase, a GTP binding protein and a transcription factor, NusA. In ModP2 OFF, proteins showing increased expression included the major RTX toxin, ApxIIIA, as well as a different set of proteins involved in general metabolism compared to ModP2 ON, such as glucose-6-phosphate 1-dehydrogenase, 2-dehydro-3- deoxyphosphogluconate aldolase and several amino acid biosynthesis proteins. The expression of several CRISPR associated proteins, three stress response factors (a large-conductance mechanosensitive channel, an excinuclease ABC subunit A, and sigma-factor 32) were also upregulated in ModP2 OFF, as were several proteins involved in fatty acid and phospholipid metabolism (putative acyl CoA thioester hydrolase, long-chain-fatty-acid--CoA ligase). ModP2 OFF also resulted in increased expression of key outer-membrane components, including a galactosyltransferase involved in lipooligosaccharide biosynthesis.

**Table 4.** Differentially regulated proteins (≥2.0 fold) in the ModP2 phasevarion.

| GenPept accession no. | Protein | Fold change<sup>a</sup> | P |
| --- | --- | --- | --- |
| **Upregulated in *modP2* ON** |  |  |  |
| lcl\|CP000687.1_prot_ABY6974<br>6.1_1140 | 30S ribosomal protein S18 | 2.73 | 0 |
| lcl\|CP000687.1_prot_ABY6987<br>2.1_1266 | cell division protein | 2.47 | 0.000118073 |
| lcl\|CP000687.1_prot_ABY6919<br>9.1_593 | transcription termination-antitermination factor, N utilization substance protein A | 2.22 | 3.79696E-14 |
| lcl\|CP000687.1_prot_ABY6916<br>4.1_558 | 50S ribosomal protein L13 | 2.17 | 0 |
| lcl\|CP000687.1_prot_ABY6910<br>9.1_503 | GTP-binding protein | 2.10 | 0.003240173 |
| lcl\|CP000687.1_prot_ABY7034<br>5.1_1739 | ribosomal protein L3 | 2.09 | 0 |
| lcl\|CP000687.1_prot_ABY7045<br>7.1_1851 | probable biotin carboxyl carrier protein of acetyl-CoA carboxylase | 2.08 | 2.22045E-16 |
| lcl\|CP000687.1_prot_ABY6998<br>0.1_1374 | 30S ribosomal protein S12 | 2.06 | 3.0332E-08 |
| lcl\|CP000687.1_prot_ABY7013<br>8.1_1532 | 30S ribosomal protein S20 | 2.00 | 4.33653E-13 |
| **Downregulated in *modP2* ON** |  |  |  |
| lcl\|CP000687.1_prot_ABY6882<br>3.1_217 | CRISPR-associated protein | 5.81 | 3.43312E-06 |
| lcl\|CP000687.1_prot_ABY6882<br>2.1_216 | CRISPR-associated protein | 4.82 | 1.82401E-05 |
| lcl\|CP000687.1_prot_ABY6988<br>0.1_1274 | glucose-6-phosphate 1-dehydrogenase | 3.38 | 9.47071E-08 |
| lcl\|CP000687.1_prot_ABY6954<br>8.1_942 | putative acyl CoA thioester hydrolase | 3.36 | 1.66756E-07 |
| lcl\|CP000687.1_prot_ABY7042<br>2.1_1816 | RNA polymerase sigma-32 factor | 3.29 | 2.28241E-05 |
| lcl\|CP000687.1_prot_ABY6994<br>6.1_1340 | cytochrome c-type biogenesis ATP-binding protein | 3.17 | 1.71531E-05 |
| lcl\|CP000687.1_prot_ABY7017<br>7.1_1571 | large-conductance mechanosensitive channel | 2.95 | 1.66276E-06 |
| lcl\|CP000687.1_prot_ABY6985<br>7.1_1251 | putative anti-sigma factor B antagonist | 2.80 | 2.79039E-06 |
| lcl\|CP000687.1_prot_ABY6990<br>2.1_1296 | RTX-III toxin determinant A from serotype 8 | 2.68 | 2.00981E-08 |
| Icl\|CP000687.1_prot_ABY6955<br>8.1_952 | lipooligosaccharide<br>galactosyltransferase I | 2.55 | 2.57572E-14 |
| Icl\|CP000687.1_prot_ABY6934<br>9.1_743 | excinuclease ABC subunit A | 2.54 | 3.19712E-08 |
| Icl\|CP000687.1_prot_ABY7036<br>5.1_1759 | preprotein translocase SecY<br>subunit | 2.49 | 0 |
| Icl\|CP000687.1_prot_ABY6938<br>4.1_778 | purine nucleotide synthesis<br>repressor | 2.44 | 3.37474E-07 |
| Icl\|CP000687.1_prot_ABY6873<br>8.1_132 | HTH-type transcriptional<br>regulator | 2.43 | 3.02394E-07 |
| Icl\|CP000687.1_prot_ABY6893<br>4.1_328 | selenide, water dikinase | 2.37 | 0.001943<br>867 |
| Icl\|CP000687.1_prot_ABY6900<br>7.1_401 | long-chain-fatty-acid--CoA ligase | 2.27 | 1.42109E-14 |
| Icl\|CP000687.1_prot_ABY6926<br>3.1_657 | DNA topoisomerase I | 2.24 | 0 |
| Icl\|CP000687.1_prot_ABY7011<br>6.1_1510 | ribosomal protein L11<br>methyltransferase | 2.18 | 5.69989E-13 |
| Icl\|CP000687.1_prot_ABY6984<br>7.1_1241 | ATP-dependent Clp protease,<br>ATP-binding subunit | 2.18 | 0 |
| Icl\|CP000687.1_prot_ABY7038<br>1.1_1775 | BC-type Fe<sup>3+</sup>-hydroxamate<br>transport system, periplasmic<br>component | 2.15 | 8.87205E-08 |
| Icl\|CP000687.1_prot_ABY7013<br>9.1_1533 | TRK system potassium uptake<br>protein | 2.14 | 0 |
| Icl\|CP000687.1_prot_ABY6877<br>2.1_166 | putative iron-sulfur binding<br>NADH dehydrogenase | 2.14 | 3.53401E-06 |
| Icl\|CP000687.1_prot_ABY6960<br>7.1_1001 | prephenate dehydratase /<br>chorismate mutase | 2.11 | 5.7693E-09 |
| Icl\|CP000687.1_prot_ABY6959<br>2.1_986 | 2-dehydro-3-deoxyphosphogluconate<br>aldolase | 2.10 | 1.44871E-11 |
| Icl\|CP000687.1_prot_ABY7008<br>9.1_1483 | serine acetyltransferase | 2.07 | 1.37564E-06 |
| Icl\|CP000687.1_prot_ABY6864<br>9.1_43 | putative tRNA/rRNA<br>methyltransferase | 2.02 | 1.74394E-11 |

<sup>a</sup> Fold change presented as *modP2* ON versus *modP2* OFF.

### ModQ is a phase-variable adenine specific DNA methyltransferase that controls a phasevarion

We also characterised the second observed *mod* gene, *modQ* that contained a GCACA_(n)_ repeat tract in *A. pleuropneumoniae*, (23) (Figure 5A). In order to determine if the *modQ* gene was also phase-variable, we also isolated a triplet set of strains with three consecutive repeat tract lengths in strain JL03 (GCACA_(n)_ repeat tracts of 9, 10, and 11 repeats; Figure 5B). Using ModQ specific antisera, we demonstrated that ModQ was only expressed when there were 9 GCACA repeats present in the SSR tract (Figure 5C, full blot in Supplementary Figure S8), and was OFF (not expressed) when either 10 or 11 GCACA repeats were present (Figure 5C). As our prototype strain JL03 encoded both ModQ and the ModP2 allele, we carried out Western blotting with our ModP2 and ModQ triplet set of strains to ensure that the *mod* gene under study contained the second *mod* gene in the same ON-OFF state, i.e, did our *modQ* ON-OFF pair contain the *modP2* gene in the same state, and vice-versa. These Western blots (Supplementary Figure S9) demonstrated that in our ModQ ON-OFF triplet, where ModQ is either 9 GCACA repeats ON, or 10 or 11 GCACA repeats OFF (anti-ModQ antisera), *modP2* was OFF in all three strains (anti-ModP sera with the same ModQ cell lysates). Analysis of the ModP2 triplet strain set showed that ModQ was ON in the ModP2 strain where there were 23 GCACA repeats, but ModQ was OFF in strains encoding ModP2 25 repeats ON, or ModP2 24 repeats OFF (anti-ModQ sera with the same ModP2 cell lysates). Therefore, for phenotypic analysis, we would use ModP2 ON GCACA 25 repeats, and ModP2 OFF GCACA 24 repeats (as described above).

**Figure 5.**
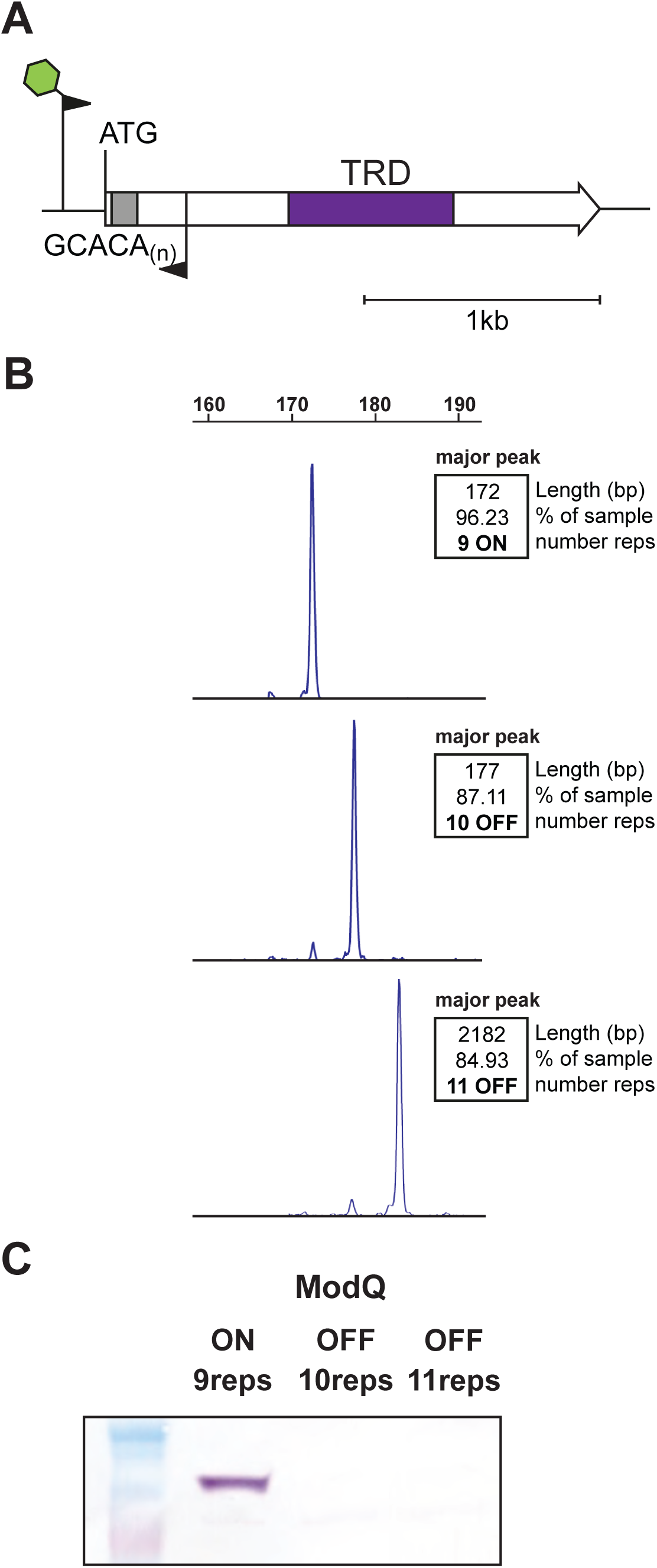
Phase-variable expression of *modQ*. (A) Illustration of the Type III *modQ* gene present in *A. pleuropneumoniae*. The *modQ* gene is present as single allelic variant (a single TRD, represented by the purple box), and like modP, also contains a variable number of GCACA_(n)_ repeats immediately downstream of the ATG start codon. The green star on the forward primer depicts a 6-Carboxyfluoresceine (FAM) fluorescent label. (B) Fragment length analysis PCR over the GCACA_(n)_ repeat tract from enriched *modQ* strains with three consecutive SSR tract lengths in strain JL03; (C) Western blot analysis using anti-ModQ sera confirmed phase-variable expression of ModQ, which was only present in a population of *A. pleuropneumoniae* enriched for 9 repeats.

Methylome analysis using genomic DNA from our triplet set of *A. pleuropneumoniae* JL03 strains encoding ModQ ON or OFF, and plasmid and genomic DNA from over-expression of ModQ and *modQ* knockout *A. pleuropneumoniae* mutant strain, demonstrated that the ModQ methyltransferase was an adenine specific Type III DNA methyltransferase, recognising the motif AG**^(m6)^A**TG (Table 2; Supplementary Data 2). This was confirmed using both PacBio SMRT sequencing and Oxford Nanopore long read sequencing and methylome analysis (Table 2; Supplementary Data 2).

Quantitative SWATH proteomics with a ModQ ON-OFF pair (9 repeats ON and 10 repeats OFF) demonstrated a total of 86 proteins had an expression level of ≥2.0 fold difference between ModQ ON-OFF pair, with 62 proteins upregulated and 24 proteins downregulated in expression in ModQ ON strain (Table 5). In ModQ ON strain, several proteins were upregulated including major RTX toxin, ApxIIIC, outer membrane lipoprotein VacJ, cell envelop proteins and several stress response proteins such as excinuclease ABC subunit A (UvrA), conserved glutaredoxin-like protein, peptide methionine sulfoxide reductase, thioredoxin (TrxA) and survival protein E (SurE). Proteins involved in metabolic pathway as well as amino acid biosynthesis proteins were also showed higher expression in ModQ ON strain. A DNA biosynthetic enzyme, as well as proteins involved in nutrient uptake and energy metabolism showed increased expression in ModQ ON strain compared to ModQ OFF strain. In the ModQ OFF strain, several ribosomal proteins, and proteins involved in fatty acid and phospholipid metabolism (probable biotin carboxyl carrier protein of acetyl-CoA carboxylase, *accB*) were all upregulated. Three major outer membrane proteins were also upregulated in ModQ OFF strain compared to ModQ ON strain (OmpH family outer membrane protein, MomP1 and ModP2). Two cell division proteins (ZipA and FtsZ) also showed increased expression in ModQ OFF compared to ModQ ON strain (Table 5). In addition, examination of OMP fractions from ModQ ON and ModQ OFF showed gross differences in outer-membrane protein expression (Supplementary Figure S7).

**Table 5.**
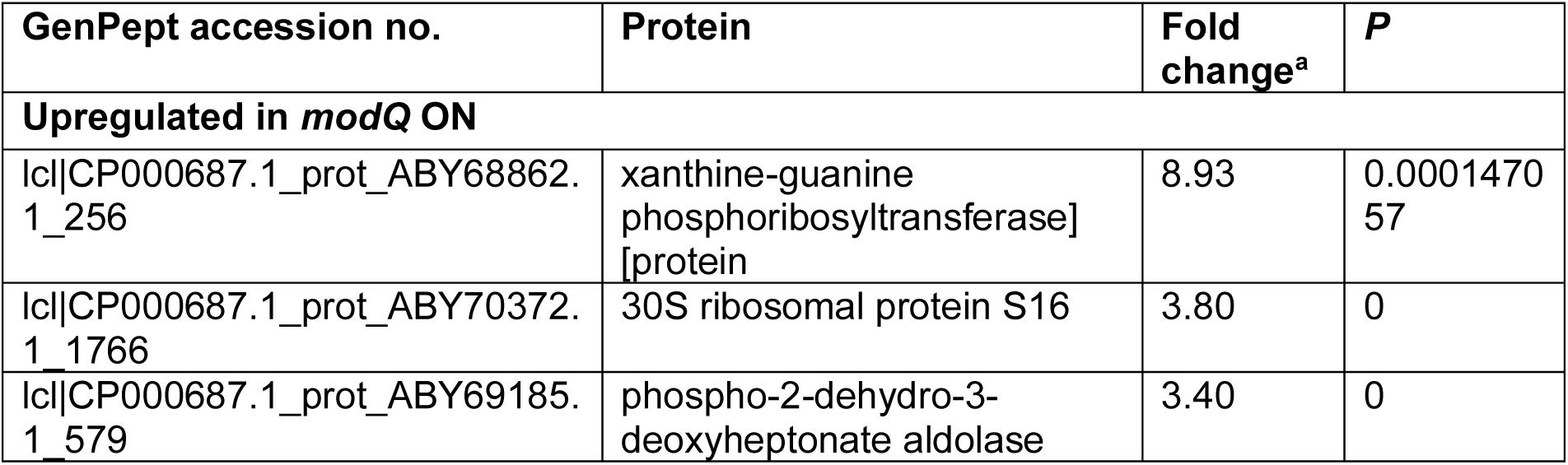
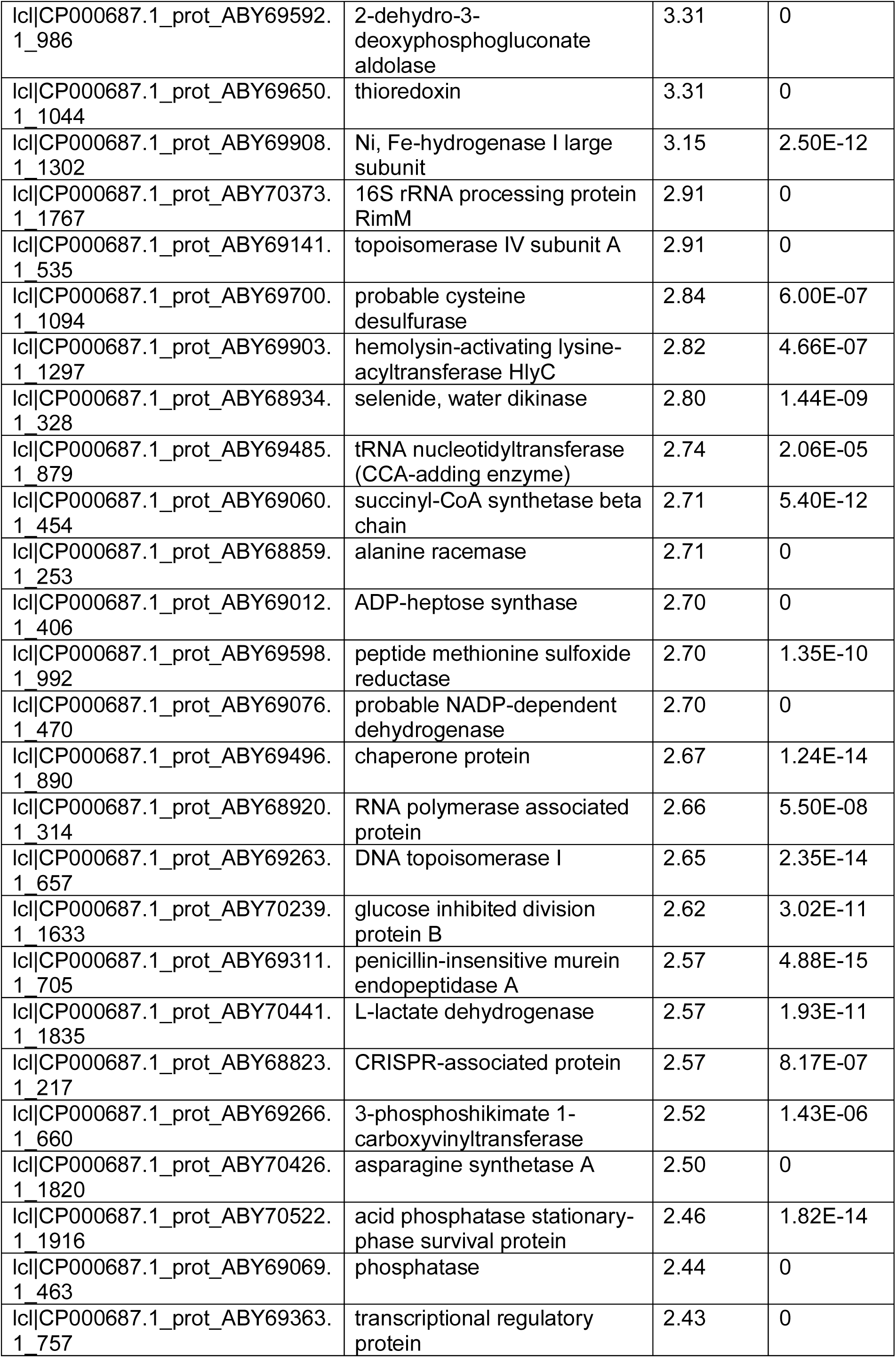
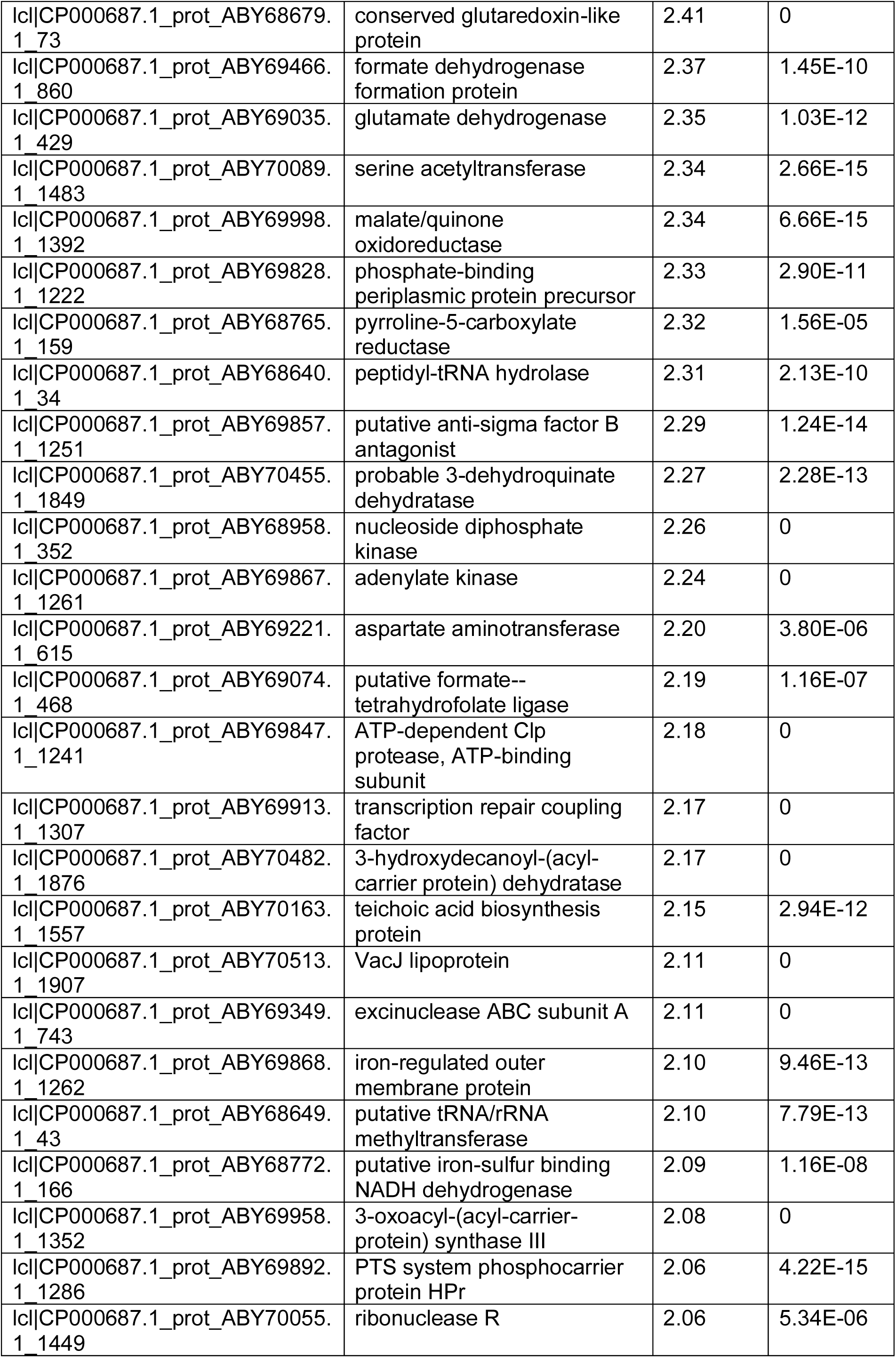
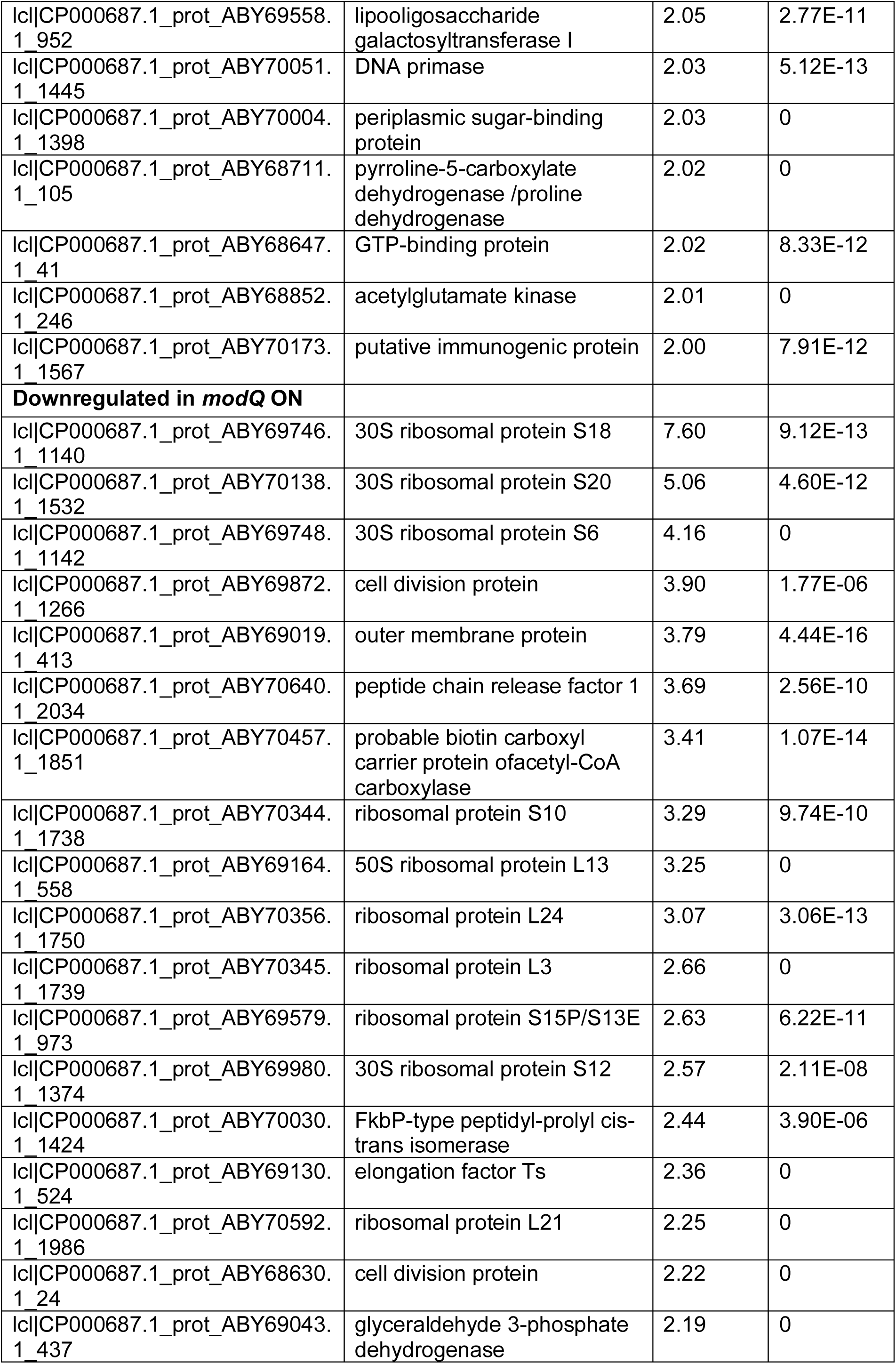
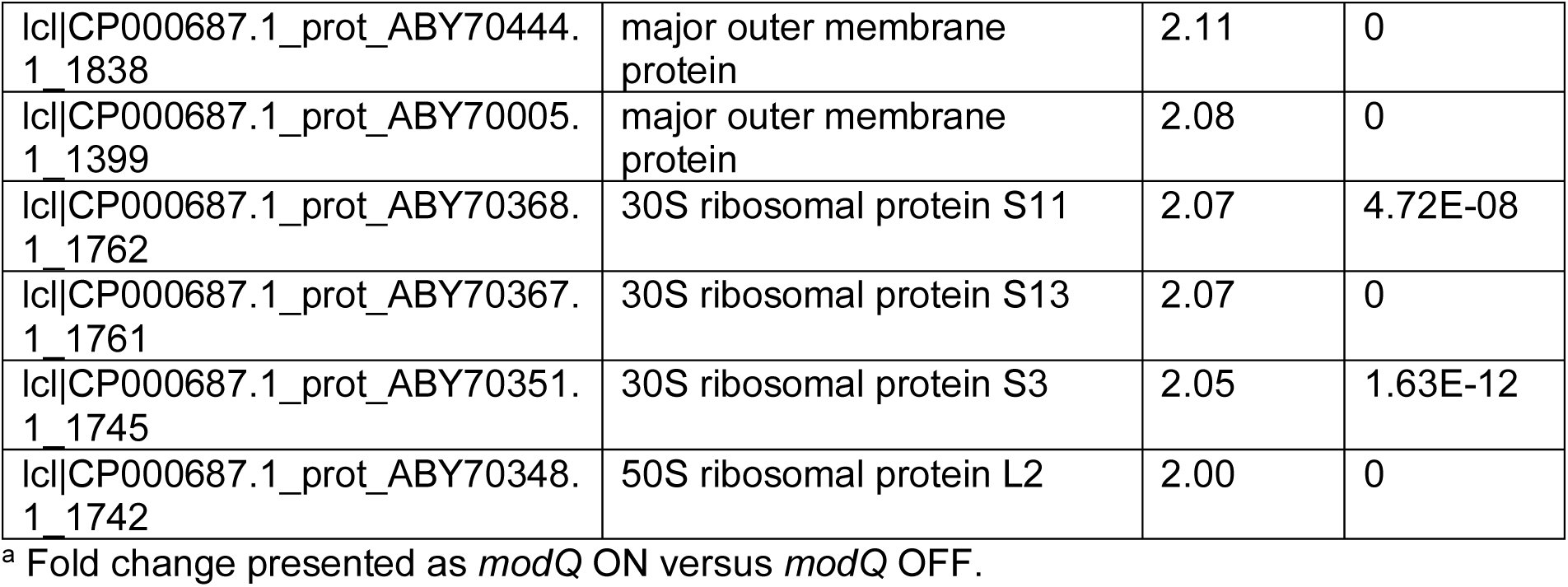
Differentially regulated proteins (≥2.0 fold) in the ModQ phasevarion.

### The ModP1 phasevarion regulates differences in growth rate and biofilm formation

We conducted standard growth curves of our enriched *mod* ON/OFF variants (*modP1*, *modP2* and *modQ*) to determine if switching of *mod* genes could affect growth rates of *A. pleuropneumoniae*. Strain AP76 expressing ModP1 (ModP1 ON) exhibited a significantly higher growth rate compared to its isogenic strain that did not express *modP1* (ModP1 OFF) (Figure 6A). Phase-variable expression of both *modP2* and *modQ* had no impact on growth rate (Figure 6B and 6C). A standard static assay biofilm assay found that biofilms formed by the *modP1* ON variant were larger in mass than its isogenic *modP1* OFF variant (Figure 6A). No differences in biofilm formation were found between either the ModP2 or the ModQ ON vs OFF variants (Figure 6B and 6C), indicating that both growth rate and biofilm formation are specifically regulated by ModP1.

**Figure 6.**
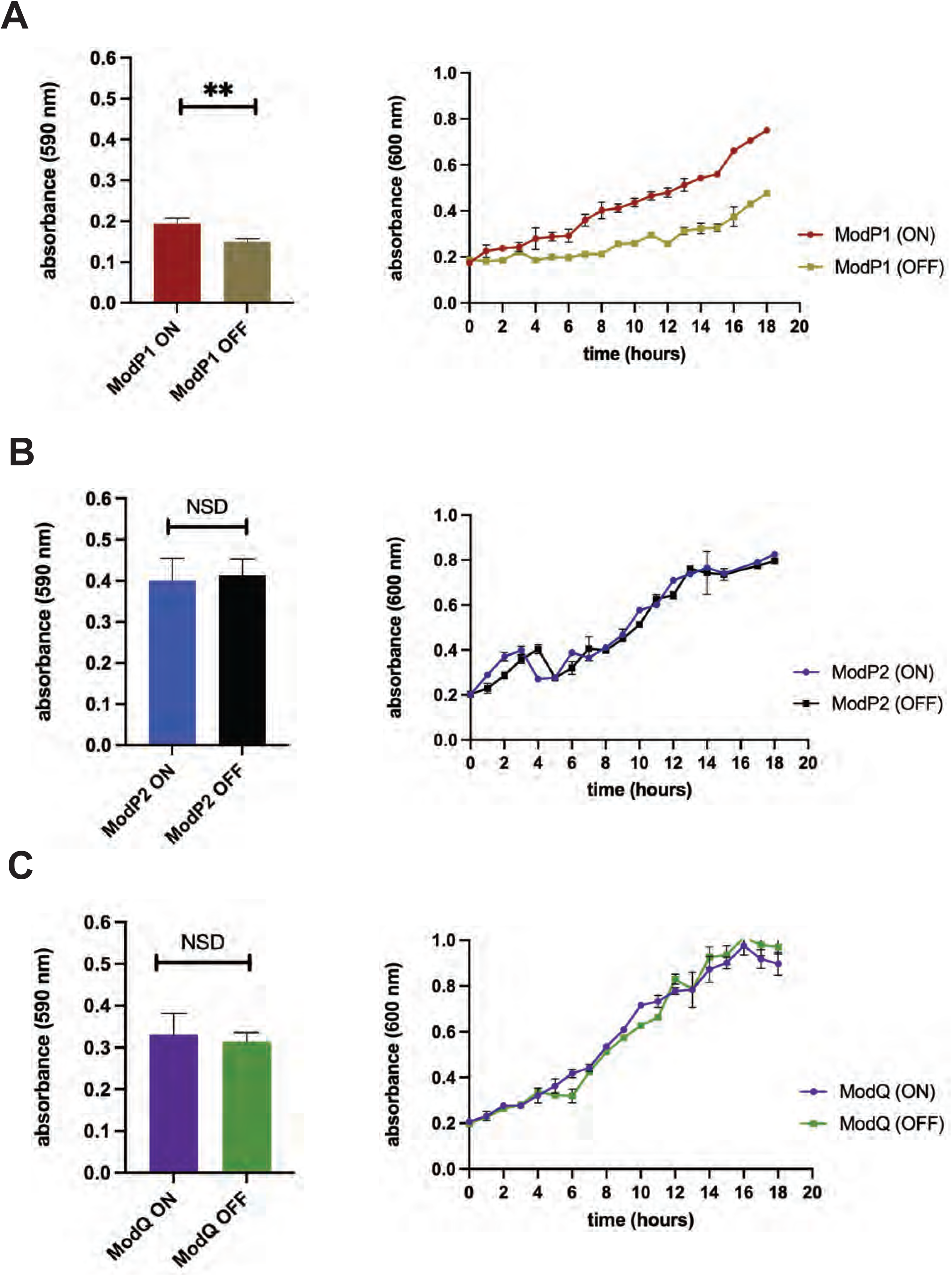
The impact of ModP and ModQ phase-variation on growth rate and biofilm formation was investigated with enriched ModP and ModQ ON/OFF variants. (A) The ModP1 ON variant showed a higher growth rate and formed a statistically larger biofilm (determined with Student’s t-test; ** - *P* = <0.01) than those that did not express ModP1 (OFF). (B) and (C) Phase-variable expression of ModP2 an ModQ had no impact on growth rate and biofilm formation (no significant difference, NSD; assessed using Student’s t-test).

### Phase variation of ModP2 influences susceptibility to antibiotics

Some human-adapted pathogens showed differential susceptibility to antibiotics due to phase variation of *mod* genes (28, 68). Therefore, we aimed to demonstrate if any of the three phasevarions (ModP1, ModP2 or ModQ) influenced susceptibility to antibiotics regularly used to treat *A. pleuropneumoniae* infections in pigs. MIC is defined as the lowest concentration of antibiotic used to inhibit bacterial growth. MIC assays of our enriched ON/OFF pairs of three *mod* genes (*modP1*, *modP2* and *modQ*) were carried out to investigate susceptibility to ampicillin, penicillin, florfenicol, ceftiofur, tulathromycin, tilmicosin and tiamulin antibiotics. This analysis showed that whilst the ModP1 and ModQ phasevarions did not result in any differential responses to the antibiotics tested, expression of ModP2 (ON) resulted in significantly higher resistance (higher MIC) to all tested antibiotics except enrofloxacin compared to ModP2 OFF (Table 6). These findings suggested that global DNA methylation by ModP2 phasevarion has an impact on antibiotic susceptibility by as yet unidentified mechanisms.

**Table 6.** MICs of seven antibiotics for *A. pleuropneumoniae* strains encoding *modP1*, *modP2* and *modQ* alleles

| MIC (ug/ml) |  |  |  |  |  |  |
| --- | --- | --- | --- | --- | --- | --- |
| antibiotics (breakpoints) | ModP1 ON | ModP1 OFF | ModP2 ON | ModP2 OFF | ModQ ON | ModQ OFF |
| Ampicillin (S≤0.5; R>2) | 5 | 5 | 5 | 0.6 | 0.6 | 0.6 |
| Penicillin (S≤0.5; R>2) | 5 | 5 | 5 | 0.6 | 0.6 | 0.6 |
| Florfenicol (S≤2; R≥8) | 0.32 | 0.32 | 5 | 0.32 | 0.32 | 0.32 |
| Tilmicosin (S≤16; R≥32) | 32 | 32 | 64 | 16 | 16 | 16 |
| Tiamulin (S≤16; R≥32) | 16 | 16 | 128 | 8 | 16 | 8 |
| Enrofloxacin (S≤0.25; R>1) | 0.03 | 0.03 | 0.03 | 0.03 | 0.03 | 0.03 |
| Tulathromycin (S≤64) | 16 | 16 | 64 | 16 | 16 | 16 |

## DISCUSSION

Many host-adapted bacterial pathogens contain phase-variable genes that are typically associated with bacterial surface structures (13,69–72) and allow bacteria to randomly switch gene expression to generate phenotypically diverse bacterial population (18). Phasevarions add a further level of complexity to understanding bacterial gene regulation, as these systems randomly switch expression of multiple genes. The only way to characterise phasevarions, as the genes regulated do not contain any identifiable features, is proteomic or gene expression profile analysis in organisms encoding a phase-variable DNA methyltransferase in ON or OFF states, or expressing each of the alternate specificity proteins.

Our phylogenetic analysis reveals that a phase-variable Type I methyltransferase is present in almost all strains of *A. pleuropneumoniae,* and that this system is highly variable: different strains contain multiple variable *hsdS* loci. Sequence analysis of the *hsdS* genes present in multiple strains encoding this locus showed a high level of variability across these strains. This demonstrates that different strains of *A. pleuropneumoniae* encoding the phase-variable Type I locus encode different HsdS variants, and therefore control different phasevarions. This is in contrast to the other characterised phase-variable Type I methyltransferases in *S. pneumoniae* (SpnIII system) (34) and *S. suis* (39), where all strains contain the same HsdS variants. Our demonstration using semi-quantitative RT-qPCR showed the expression of all four HsdS variants in strains AP76 (variants A-D) and JL03 (D-G), providing good evidence that all HsdS variants are expressed and functional and that this Type I system is phase-variable. We also demonstrated that in addition to the major expressed *hsdS* allele (allele A in strain AP76 and allele E in strain JL03), minor variants do exist in populations of strain AP76 and JL03 using our bespoke FAM-labelled PCR assay. Whether this Type I system controls different phasevarions remains to be elucidated.

Our study provided a major focus on the phase-variable Type III methyltransferases encoded in *A. pleuropneumoniae.* This demonstrated the presence of two distinct phase-variable *mod* genes in *A. pleuropneumoniae* (23), which we have named *modP* and *modQ*. By analysing multiple strain collections, we identified four allelic variants of *modP* (encoding proteins ModP1-4), and a single variant of *modQ*. These *mod* genes show a distinct conservation within different serovars of *A. pleuropneumoniae*. We confirmed that both *modP* and *modQ* switch expression ON-OFF due to changes in length of a locus encoded GCACA_(n)_ SSR tract using prototype strains encoding *modP1* (strain AP76), *modP2* (strain JL03) and *modQ* (strain JL03) (Figure 4 and 5) using our well-established FAM-labelled PCR coupled to fragment length analysis (28,39,40,73). Western blotting using antisera against the conserved region of ModP and ModQ confirmed expression of each gene is commensurate with GCACA_(n)_ SSR tract length, confirming that all these Mod proteins are phase-variable.

Using enriched ON-OFF pairs in our prototype strains for ModP1, ModP2, and ModQ, we used a combination of SMRT sequencing and Oxford Nanopore sequencing with technology specific methylome analysis to determine the specificity of each Mod protein. Although PacBio long read technology is well established as the gold-standard to solve methyltransferase specificities *de novo* (74, 75), Nanopore sequencing to solve the specificity of bacterial DNA methyltransferases has also been used effectively (55, 74). This analysis demonstrated that ModQ is an adenine specific phase-variable DNA methyltransferase, methylating the sequence 5′-AG**^(m6)^A**TG-3′. All other Type III Mod proteins that have been shown to control phasevarions are also adenine specific DNA methyltransferases (28,39,40,73). Our methylome analysis demonstrated that ModP is cytosine-specific, making it the first identified and characterised phase-variable cytosine-specific Mod that controls a phasevarion. In common with all other Mod proteins, different allelic variants of ModP methylate different DNA sequences (ModP1 methylates 5′-CAA**^(m4)^C**T-3′ and ModP2 methylates 5′- GW**^(m4)^C**CT-3′). Whilst we used the gold standard of PacBio SMRT sequencing and methylome analysis to conclusively show that ModP1 recognises 5′-CAA**^(m4)^C**T-3′, we used a combination of PacBio and Nanopore methodologies to determine the ModP2 specificity of 5′-GW**^(m4)^C**CT-3′. Both were shown to be cytosine specific, with different specificities due to each allele encoding a different TRD. Interestingly we could not detect methylation by ModP1 using Nanopore sequencing of genomic DNA isolated from either *E. coli* over-expressing this methyltransferase, or from *A. pleuropneumoniae* strains encoding ModP1. We were also unable to detect ModP2 methylation using *A. pleuropneumoniae* genomic DNA. Despite rigorous analysis and refinement of our methodology, it remains unclear why this is the case, and beyond the scope of this current work, as adenine methylation by ModQ was easily detected using Nanopore technology with all DNA used, confirming our findings with our PacBio methodology. Never-the-less, we confirmed methyltransferase specificity of ModP1, ModP2, and ModQ using multiple methods, conclusively demonstrating each recognises and methylates a unique DNA target sequence.

Using SWATH proteomics, a well-established method to study protein expression differences mediated by phase-variable DNA methyltransferases (40,76,77), we demonstrate that all three Mod variants (ModP1, ModP2, and ModQ) control distinct sets of proteins, i.e., they all control different phasevarions. Several proteins with key roles in *A. pleuropneumoniae* pathobiology, are all controlled by different phasevarions. These proteins include the major RTX toxins (ApxIVA, ApxIIIA and ApxIIIC) (78–80), several outer membrane proteins (80) and proteins involved in LOS and capsule biosynthesis (81), implying a complex mode of regulation of these factors, and raising questions about their suitability for use in vaccines against *A. pleuropneumoniae*. Inclusion of phase-variable proteins as components of subunit vaccines has previously been discounted, as variable expression of vaccine targets could lead to a decrease in vaccine efficacy. As we identify several proteins in multiple phasevarions that are putative *A. pleuropneumoniae* vaccine candidates, including the RTX toxins, it is critical to understand the exact mode of regulation of these proteins in order to direct and inform vaccine subunit vaccine development.

Phenotypic analysis of using our enriched ON-OFF pairs demonstrates a complex effect on the phenotype of strains encoding the phase-variable Mod proteins ModP1, ModP2, and ModQ. We reveal that ModP1 ON-OFF switching results in key differences in growth rate and biofilm formation. This variation in growth rate may give one population a selective advantage during long-term carriage. As biofilm formation is a key pathology of *A. pleuropneumoniae* (82, 83), strains encoding ModP1 could result in different disease outcomes, or even niche specific differences, dependent on the ON-OFF status of ModP1. Biofilm formation also increases resistance to antimicrobials (84–86), although we did not observe a difference in MIC with our ModP1 ON-OFF pair. Phase-variable switching of ModP2 did, however, have a major effect on the MIC of *A. pleuropneumoniae*. When ModP2 was ON, we observed significantly greater MIC using multiple classes of antibiotic when compared to the ModP2 OFF strain. This suggests that ModP2 controls expression of proteins required for antibiotic resistance and could have major effects on topical treatment of *A. pleuropneumoniae* infections in pigs, as well as implications for the spread of antibiotic resistance. For example, treating diseased pigs with standard concentrations of antibiotics may lead to a selection for the ModP2 ON state in those strains encoding this Mod allele, and lead to an emergence of resistance. Differences in antibiotic resistance with phase-variable genes (13) and other phasevarions (28) has also been observed previously.

In summary, this study demonstrates the presence of multiple phase-variable DNA methyltransferases in *A. pleuropneumoniae*, and has characterised the first ever phasevarions controlled by multiple variants of a cytosine specific phase-variable Mod, ModP. Multiple phase-variable Type III methyltransferases in *A. pleuropneumoniae* control distinct phasevarions, each of which has a unique effect on protein expression, and on phenotypes such as biofilm formation (ModP1) and antibiotic resistance (ModP2). The presence of multiple phase-variable DNA methyltransferases controlling phasevarions is likely to complicate vaccine development and topical treatment of infections using antibiotics. Defining the stably expressed antigenic repertoire of an organism is key to the rational design of subunit vaccines. As no broadly effective protective vaccine is available against *A. pleuropneumoniae*, understanding the role of phasevarions is key to not only treating infections, but also to understanding which proteins are stably expressed; by thoroughly characterising the proteins controlled by phasevarions in *A. pleuropneumoniae*, we have provided the baseline to future vaccine development against a major veterinary bacterial pathogen.

## DATA AVAILABILITY

The mass spectrometry proteomics data have been deposited to the ProteomeXchange Consortium via the PRIDE (87) partner repository with the dataset identifier PXD036273.

## FUNDING

This work was funded by an Australian Research Council Discovery Project (DP180100976) to JMA and PJB, an Australian National Health and Medical Research Council (NHMRC) Principal Research Fellowship (1138466) to MPJ, an NHMRC Investigator Grant (GNT2007996) to QG, a United Kingdom Biotechnology and Biological Research Council BBSRC grants (BB/S002103/1 and BB/G018553/1) to PRL and JB. LAW is supported by a Sir Henry Dale Fellowship jointly funded by the Wellcome Trust and the Royal Society (109385/Z/15/Z). Work at the Walter and Eliza Hall Institute of Medical Research was made possible through Victorian State Government Operational Infrastructure Support and Australian Government NHMRC IRIISS. Publication costs for this work were generously funded by the Bourne Foundation, Melbourne, Australia

## ACKNOWLEDGEMENTS

We thank Amanda Nouwens and Peter Josh in SCMB MS facility, University of Queensland for conducting SWATH-MS proteomic analysis. We thank Allison Banse and Eric Johnson at SNPsaurus (University of Oregon, USA) for conducting PacBio SMRT sequencing and methylome analysis. We thank the Australian Genome Research Facility (AGRF, Brisbane) and Nicole Hogg at the Griffith University DNA Sequencing Facility (GUDSF) for fragment length separation and analysis.

---

**Supplementary Figure S1:**
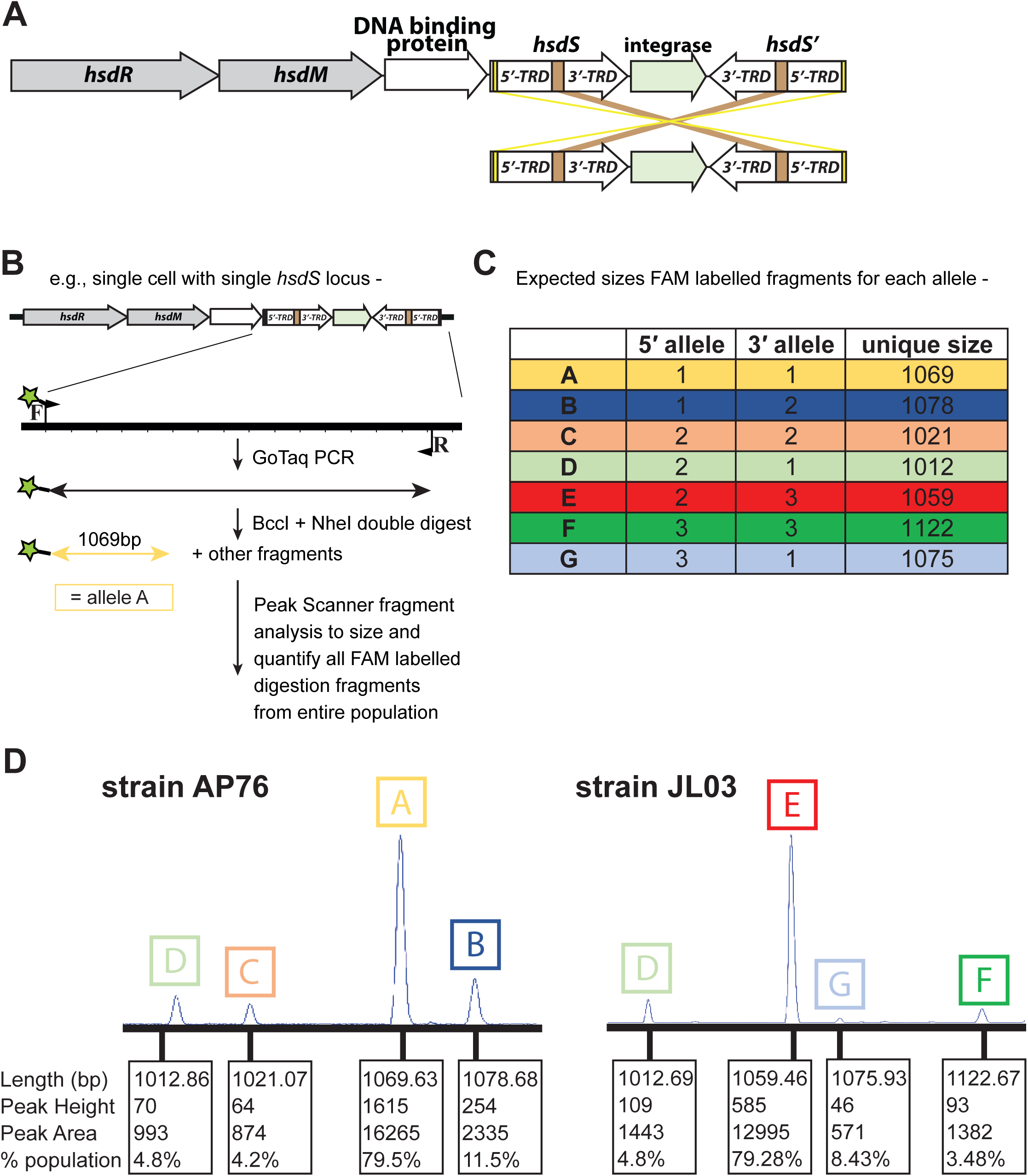
Phase-variable expression of Type I R-M system. (A) Illustration of the Type I R-M system containing duplicate, variable *hsdS* genes and IRs. Gene shuffling between variable 5’ and 3’ TRDs produces four unique *hsdS* genes at the expressed *hsdS* locus immediately downstream of the *hsdM* gene, resulting in four unique allelic variants of the HsdS protein. (B) Illustration of FAM-labelled PCR assay used to quantify *hsdS* alleles present in the expressed locus in a population of *A. pleuropneumoniae*, with FAM label illustrated as green star on forward primer. (C) predicted sizes of each allelic variant based on *in silico* analysis. (D) FAM-labelled fragments are separated and quantified using a Gene Analyser (Life Technologies) and quantified by Peak Scanner software in order to determine the percentage of each allele present in the *A. pleuropneumoniae* population.

**Supplementary Figure S2:**
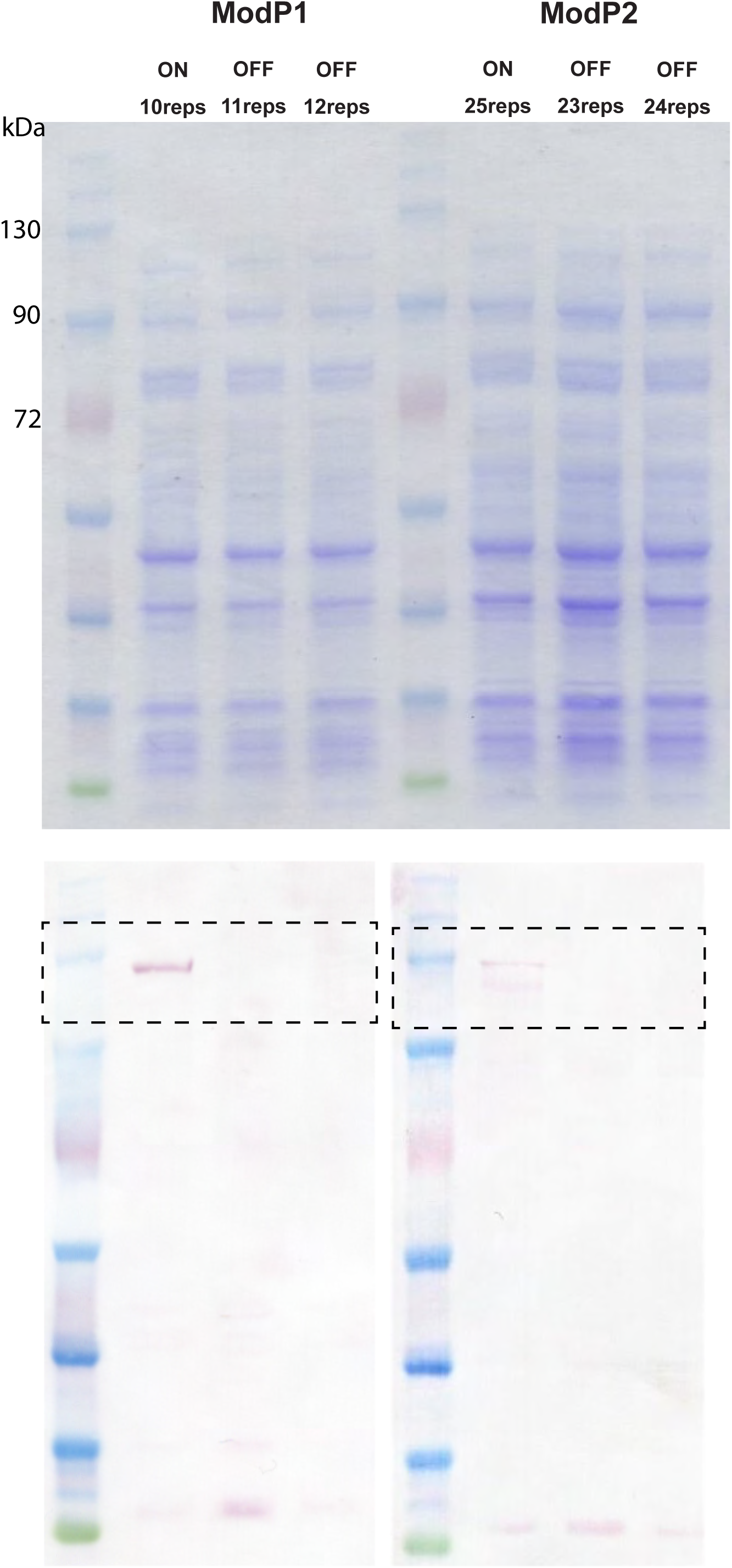
ModP Coomassie and full Western blot. Whole-celllysates from strain AP76 with 10, 11 and 12 SSR repeats and strain JL03 with 23, 24 and 25 SSR repeats were run on 4-12% Bis-Tris gel and Coomassie blue staining was carried out to visualize proteins. Western blotting with anti-ModP sera demonstrates that *modP1* gene expression is ON in a population enriched with 10 GCACA repeats and *modP2* with 25 GCACA repeats.

**Supplementary Figure S3.**
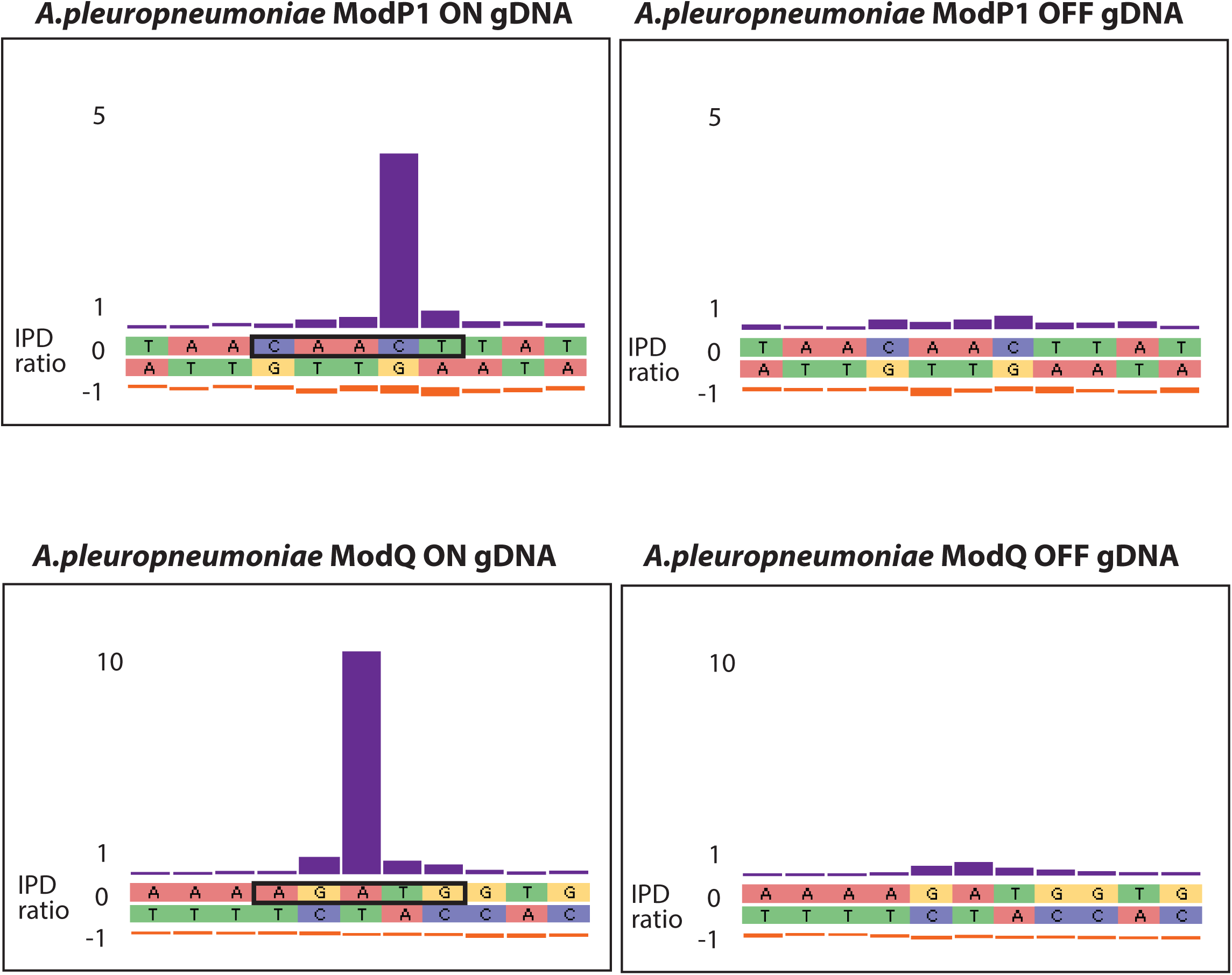
examples of motifs discovered for ModP1 and ModQ using PacBio SMRT sequencing and methylome analysis in ModP ON-OFF and ModQ ON-OFF strains. The motifs for ModP1 and ModQ are only detected when each Mod is expressed (ON).

**Supplementary Figure S4.**
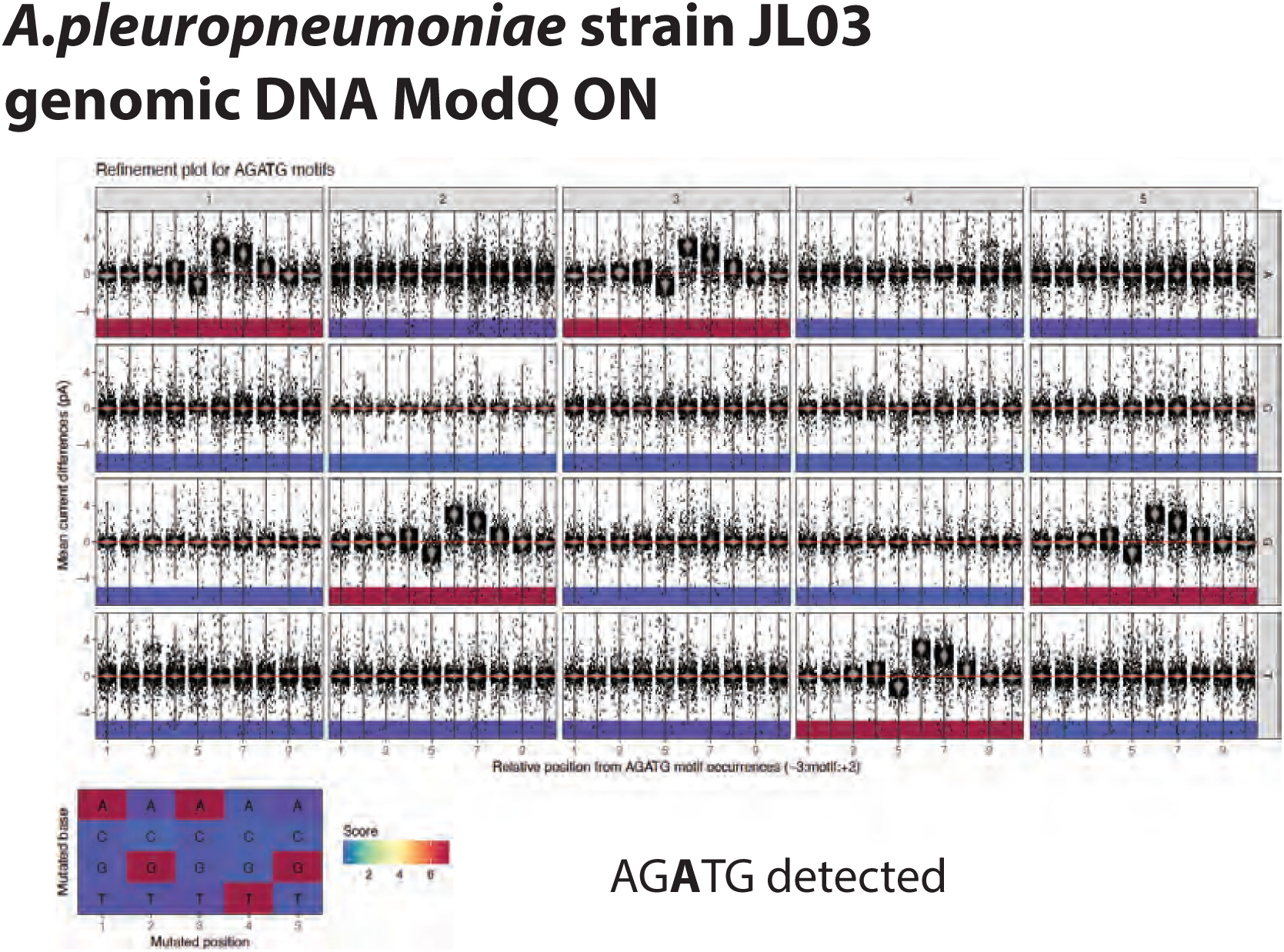
example of Nanodisco output and motifs detected using genomic DNA isolated from *A. pleuropneumoniae* ModQ ON vs ModQ::cat knockout

**Supplementary Figure S5.**
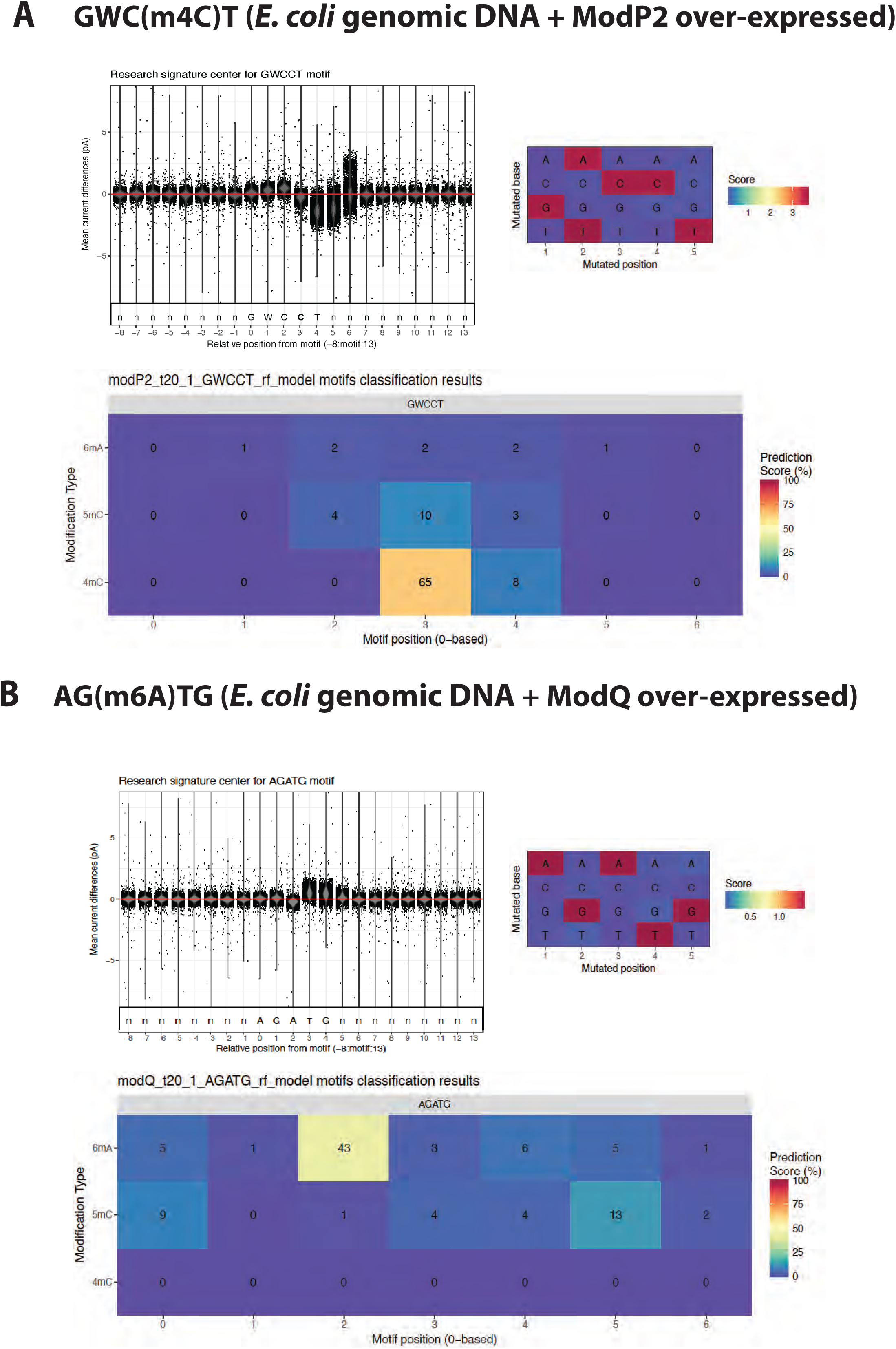
example of Nanodisco output and motifs detected using genomic DNA isolated from *E. coli* BL21 over-expressing ModP2 and ModQ when compared to *E. coli* BL21 only genomic DNA

**Supplementary Figure S6.**
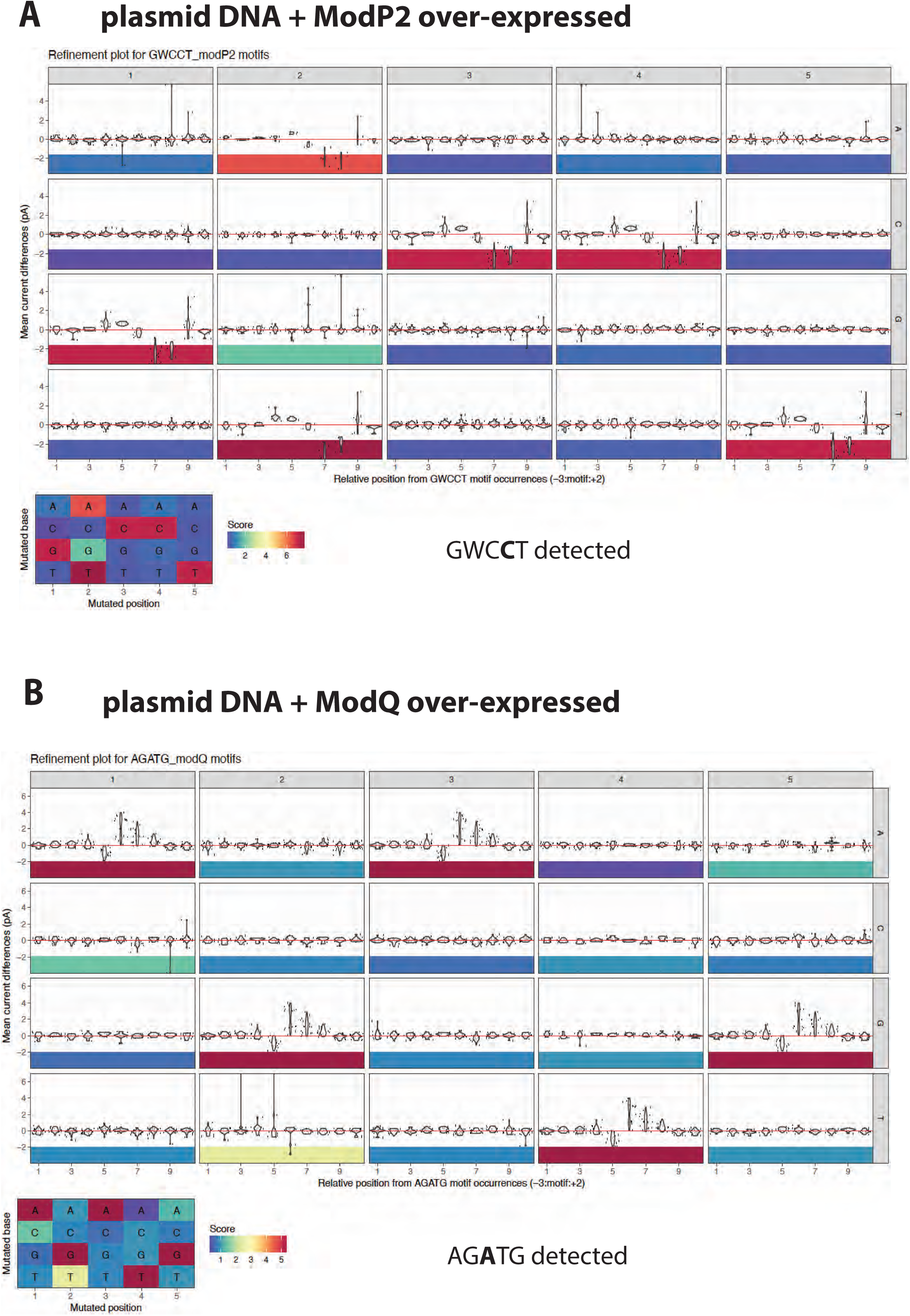
example of Nanodisco output and motifs detected using plasmid DNA isolated from *E. coli* BL21 over-expressing ModP2 and ModQ

**Supplementary Figure S7:**
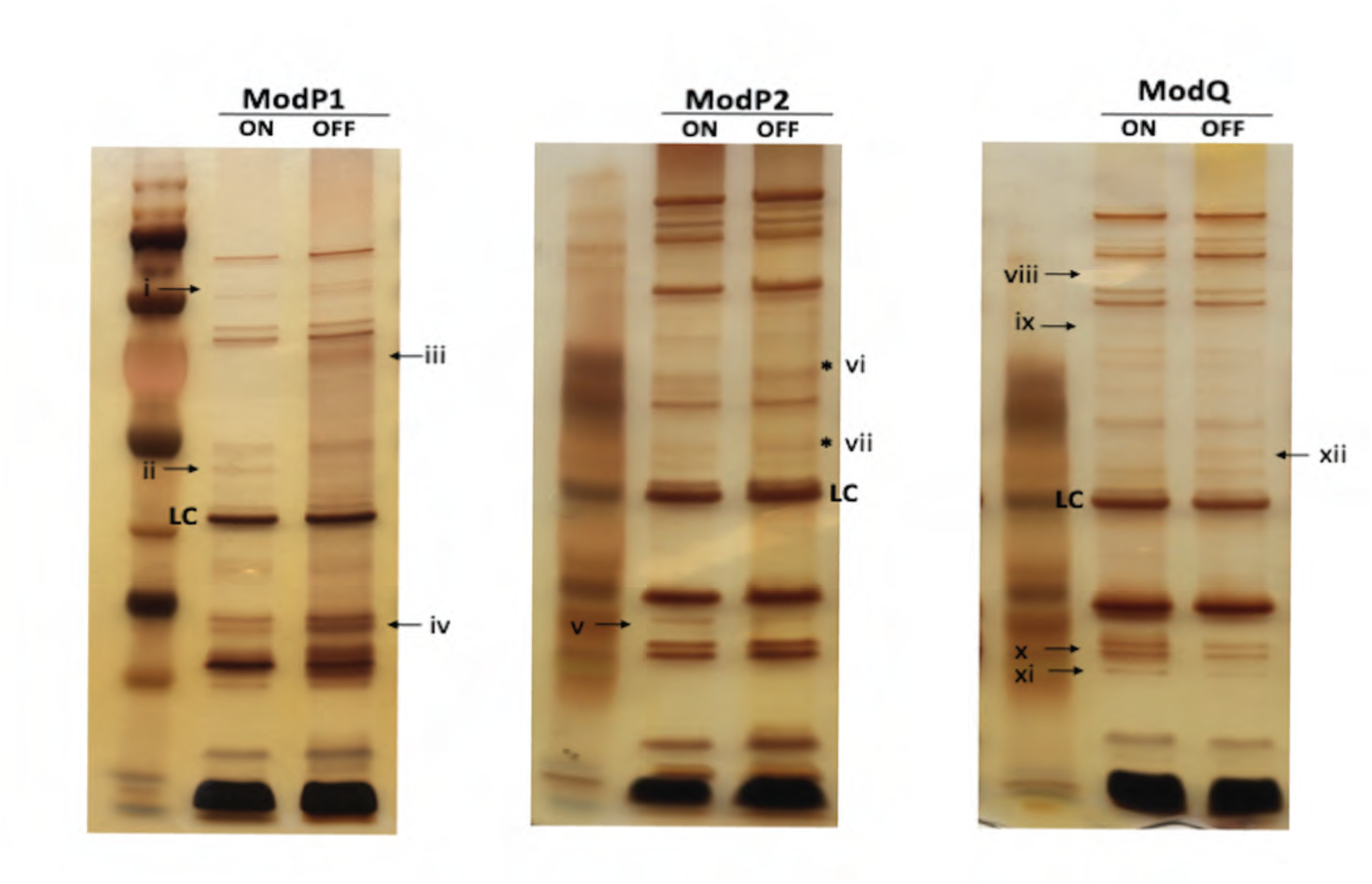
OMP silver staining. Silver-stained protein gel showing OMPs differences due to ON and OFF switching of methyltransferase genes. Differences in protein expression between ON vs OFF pairs of each mod gene are highlighted with arrows. After visually identifying any differences, each band was processed with ImageJ using a band that was observed as the same intensity in the respective ON vs OFF lane (loading control; LC). Only bands that showed a ≥2-fold intensity difference relative to the loading control are highlighted. ModP1 ON vs OFF: (i) protein band is absent in ON strain; (ii) 4 fold; (iii) protein band is absent in ON strain; (iv) 0.29 fold. ModP2 ON vs OFF: (v) 2.9 fold; (vi) and (vii) difference in size of proteins (*). ModQ ON vs OFF: (viii) and (ix) protein bands are present in ON strain; (x) 2.00 fold; (xi) 2.43 fold; (xii) protein band is absent in ON strain.

**Supplementary Figure S8:**
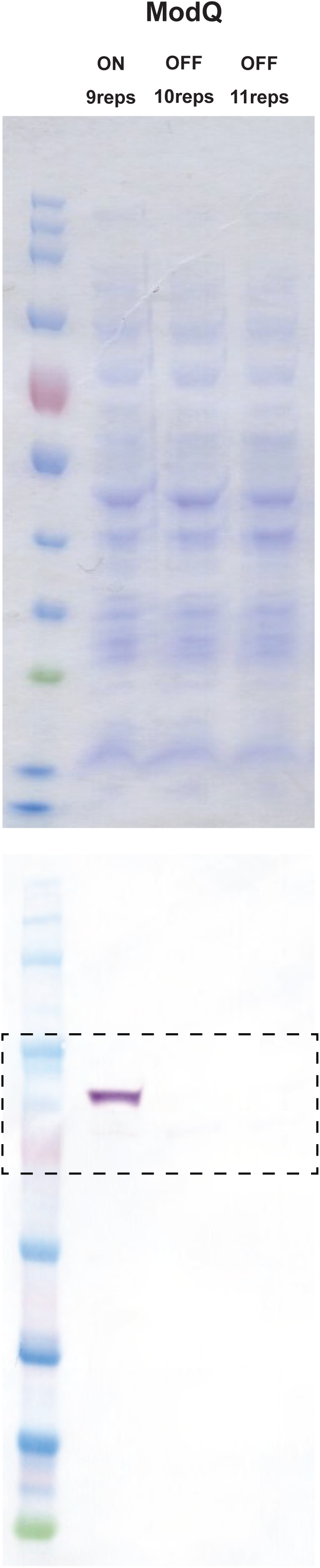
ModQ Coomassie and full blot. Whole cell lysates were prepared from strain JL03 with 9, 10 and 11 GCACA repeats and run on 4-12% Bis-Tris gel. Western blot analysis using anti-ModQ sera demonstrates that *modQ* gene expression is ON in a population enriched with 9 GCACA repeats and OFF with 10 or 11 GCACA repeats.

**Supplementary Figure S9:**
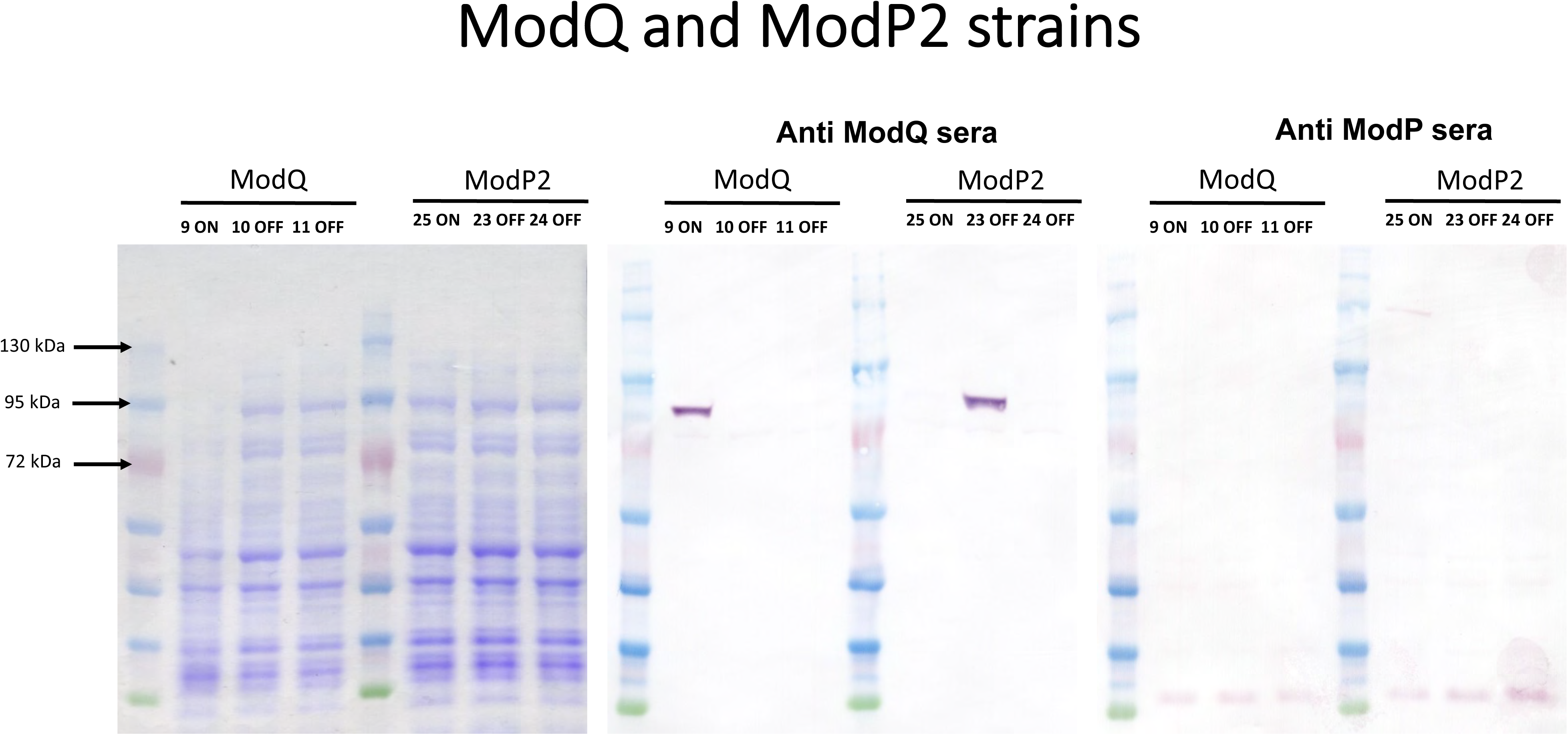
Western blots of ModP2 vs ModQ. Due to co-localisation of *modP2* and *modQ* genes in strain JL03, Western blot analysis was performed with enriched *modP2* and *modQ* ON and OFF variants using anti-ModP and anti-ModQ sera. Western blot analysis confirms that *modP2* gene expression is OFF in our ModQ ON-OFF triplet set of strains (anti-ModP sera) but the *modQ* gene was ON in the ModP2 strain where there were 23 GCACA repeats in the *modP2* gene (anti-ModQ sera).

**Supplementary Table S1.** Details of strains used to generate the phylogenetic tree In Figure 2

**Supplementary Table S2.** Details of all strains used from the Australian national reference collection for *A. pleuropneumoniae* (UQ) used to screen for Type Ill methyltransferase alleles and genes

Supplementary Table S3. Primers used in this study.

## Supplementary Data

Supplmentary Data 51 - sequence analysis of all TRD variants found in *A. pleuropneumoniae* strains with an annotated genome in REBASE

Supplementary Data 52- all methylome data from PacBio and Nanopore analysis

## Notes

### Competing Interest Statement

The authors have declared no competing interest.

## REFERENCES

1. 1. Gottschalk, M. and Broes, A. (2019) In Jeffrey J. Zimmerman, L. A. K., Alejandro Ramirez, Kent J. Schwartz, Gregory W. Stevenson, Jianqiang Zhang (ed.), Diseases of swine. 11th ed, pp. 749–766.

2. Chiers, K., Donné, E., Van Overbeke, I., Ducatelle, R. and Haesebrouck, F. (2002) *Actinobacillus pleuropneumoniae* infections in closed swine herds: infection patterns and serological profiles. Vet. Microbiol., 85, 343–352.

3. Chiers, K., Van Overbeke, I., Donné, E., Baele, M., Ducatelle, R., De Baere, T. and Haesebrouck, F. (2001) Detection of *Actinobacillus pleuropneumoniae* in cultures from nasal and tonsillar swabs of pigs by a PCR assay based on the nucleotide sequence of a dsbE-like gene. Vet. Microbiol., 83, 147–159.

4. Vigre, H., Angen, Ø., Barfod, K., Lavritsen, D.T. and Sørensen, V. (2002) Transmission of *Actinobacillus pleuropneumoniae i*n pigs under field-like conditions: emphasis on tonsillar colonisation and passively acquired colostral antibodies. Vet. Microbiol., 89, 151–159.

5. Tobias, T.J., Bouma, A., Daemen, A.J., Wagenaar, J.A., Stegeman, A. and Klinkenberg, D. (2013) Association between transmission rate and disease severity for *Actinobacillus pleuropneumoniae* infection in pigs. Vet. Res., 44, 1–10.

6. Stringer, O.W., Bossé, J.T., Lacouture, S., Gottschalk, M., Fodor, L., Angen, Ø., Velazquez, E., Penny, P., Lei, L. and Langford, P.R. (2021) Proposal of *Actinobacillus pleuropneumoniae* serovar 19, and reformulation of previous multiplex PCRs for capsule-specific typing of all known serovars. Vet. Microbiol., 255, 109021.

7. Chiers, K., van Overbeke, I., De Laender, P., Ducatelle, R., Carel, S. and Haesebrouck, F. (1998) Effects of endobronchial challenge with *Actinobacillus pleuropneumoniae* serotype 9 of pigs vaccinated with inactivated vaccines containing the Apx toxins. The Veterinary quarterly, 20, 65–69.

8. Hensel, A., Huter, V., Katinger, A., Raza, P., Strnistschie, C., Roesler, U., Brand, E. and Lubitz, W. (2000) Intramuscular immunization with genetically inactivated (ghosts) *Actinobacillus pleuropneumoniae* serotype 9 protects pigs against homologous aerosol challenge and prevents carrier state. Vaccine, 18, 2945–2955.

9. Bayliss, C.D. (2009) Determinants of phase variation rate and the fitness implications of differing rates for bacterial pathogens and commensals. FEMS Microbiol. Rev., 33, 504–520.

10. Moxon, R., Bayliss, C. and Hood, D. (2006) Bacterial contingency loci: the role of simple sequence DNA repeats in bacterial adaptation. Annu. Rev. Genet., 40, 307–333.

11. Power, P.M., Sweetman, W., Gallacher, N., Woodhall, M., Kumar, G., Moxon, E. and Hood, D. (2009) Simple sequence repeats in *Haemophilus influenzae*. *Infect.*, Genet. Evol., 9, 216–228.

12. Saunders, N.J., Jeffries, A.C., Peden, J.F., Hood, D.W., Tettelin, H., Rappuoli, R. and Moxon, E.R. (2000) Repeat-associated phase variable genes in the complete genome sequence of *Neisseria meningitidis* strain MC58. Mol. Microbiol., 37, 207–215.

13. Atack, J.M., Winter, L.E., Jurcisek, J.A., Bakaletz, L.O., Barenkamp, S.J. and Jennings, M.P. (2015) Selection and counterselection of Hia expression reveals a key role for phase-variable expression of Hia in infection caused by nontypeable *Haemophilus influenzae*. The Journal of Infectious Diseases, 212, 645–653.

14. Elango, D. and Schulz, B.L. (2019) Phase-variable glycosylation in nontypeable *Haemophilus influenzae*. Journal of Proteome Research, 19, 464–476.

15. Blyn, L.B., Braaten, B.A. and Low, D.A. (1990) Regulation of *pap* pilin phase variation by a mechanism involving differential dam methylation states. The EMBO Journal, 9, 4045–4054.

16. Tzeng, Y.-L., Thomas, J. and Stephens, D.S. (2016) Regulation of capsule in *Neisseria meningitidis*. Crit. Rev. Microbiol., 42, 759–772.

17. Phillips, Z.N., Brizuela, C., Jennison, A.V., Staples, M., Grimwood, K., Seib, K.L., Jennings, M.P. and Atack, J.M. (2019) Analysis of invasive nontypeable *Haemophilus influenzae* isolates reveals selection for the expression state of particular phase-variable lipooligosaccharide biosynthetic genes. Infect. Immun., 87, e00093–00019.

18. Moxon, R., Bayliss, C. and Hood, D. (2006) Bacterial contingency loci: the role of simple sequence DNA repeats in bacterial adaptation. Annu. Rev. Genet., 40, 307–333.

19. Atack, J.M., Tan, A., Bakaletz, L.O., Jennings, M.P. and Seib, K.L. (2018) Phasevarions of bacterial pathogens: methylomics sheds new light on old enemies. Trends Microbiol., 26, 715–726.

20. Phillips, Z.N., Husna, A.-U., Jennings, M.P., Seib, K.L. and Atack, J.M. (2019) Phasevarions of bacterial pathogens–phase-variable epigenetic regulators evolving from restriction– modification systems. Microbiology, 165, 917–928.

21. Seib, K.L., Srikhanta, Y.N., Atack, J.M. and Jennings, M.P. (2020) Epigenetic regulation of virulence and immunoevasion by phase-variable restriction-modification systems in bacterial pathogens. Annu. Rev. Microbiol., 74, 655–671.

22. Roberts, R.J., Vincze, T., Posfai, J. and Macelis, D. (2015) REBASE—a database for DNA restriction and modification: enzymes, genes and genomes. Nucleic Acids Res., 43, D298–D299.

23. Atack, J.M., Yang, Y., Seib, K.L., Zhou, Y. and Jennings, M.P. (2018) A survey of Type III restriction-modification systems reveals numerous, novel epigenetic regulators controlling phase-variable regulons; phasevarions. Nucleic Acids Res., 46, 3532–3542.

24. Atack, J.M., Guo, C., Yang, L., Zhou, Y. and Jennings, M.P. (2020) DNA sequence repeats identify numerous Type I restriction-modification systems that are potential epigenetic regulators controlling phase-variable regulons; phasevarions. The FASEB Journal, 34, 1038–1051.

25. Atack, J.M., Guo, C., Litfin, T., Yang, L., Blackall, P.J., Zhou, Y. and Jennings, M.P. (2020) Systematic analysis of REBASE identifies numerous type I restriction-modification systems with duplicated, distinct hsdS specificity genes that can switch system specificity by recombination. Msystems, 5, e00497–00420.

26. Srikhanta, Y.N., Dowideit, S.J., Edwards, J.L., Falsetta, M.L., Wu, H.-J., Harrison, O.B., Fox, K.L., Seib, K.L., Maguire, T.L. and Wang, A.H.-J. (2009) Phasevarions mediate random switching of gene expression in pathogenic Neisseria. PLoS Path., 5, e1000400.

27. Srikhanta, Y.N., Maguire, T.L., Stacey, K.J., Grimmond, S.M. and Jennings, M.P. (2005) The phasevarion: A genetic system controlling coordinated, random switching of expression of multiple genes. Proc. Natl. Acad. Sci. U. S. A., 102, 5547–5551.

28. Atack, J.M., Srikhanta, Y.N., Fox, K.L., Jurcisek, J.A., Brockman, K.L., Clark, T.A., Boitano, M., Power, P.M., Jen, F.E.-C. and McEwan, A.G. (2015) A biphasic epigenetic switch controls immunoevasion, virulence and niche adaptation in non-typeable *Haemophilus influenzae*. Nature communications, 6, 1–12.

29. Srikhanta, Y.N., Gorrell, R.J., Steen, J.A., Gawthorne, J.A., Kwok, T., Grimmond, S.M., Robins-Browne, R.M. and Jennings, M.P. (2011) Phasevarion mediated epigenetic gene regulation in *Helicobacter pylori*. PloS one, 6, e27569.

30. Blakeway, L.V., Power, P.M., Jen, F.E.C., Worboys, S.R., Boitano, M., Clark, T.A., Korlach, J., Bakaletz, L.O., Jennings, M.P. and Peak, I.R. (2014) ModM DNA methyltransferase methylome analysis reveals a potential role for *Moraxella catarrhalis* phasevarions in otitis media. The FASEB Journal, 28, 5197–5207.

31. Gawthorne, J.A., Beatson, S.A., Srikhanta, Y.N., Fox, K.L. and Jennings, M.P. (2012) Origin of the diversity in DNA recognition domains in phasevarion associated *modA* genes of pathogenic *Neisseria* and *Haemophilus influenzae*. PLoS One, 7, e32337.

32. Tan, A., Hill, D.M., Harrison, O.B., Srikhanta, Y.N., Jennings, M.P., Maiden, M.C. and Seib, K.L. (2016) Distribution of the type III DNA methyltransferases *modA*, *modB* and *modD* among *Neisseria meningitidis* genotypes: implications for gene regulation and virulence. Sci Rep, 6, 21015.

33. Gorrell, R. and Kwok, T. (2017) The *Helicobacter pylori* methylome: roles in gene regulation and virulence. Molecular Pathogenesis and Signal Transduction by Helicobacter pylori.

34. Manso, A.S., Chai, M.H., Atack, J.M., Furi, L., De Ste Croix, M., Haigh, R., Trappetti, C., Ogunniyi, A.D., Shewell, L.K., Boitano, M. et al. (2014) A random six-phase switch regulates pneumococcal virulence via global epigenetic changes. Nature communications, 5, 5055.

35. Loenen, W.A., Dryden, D.T., Raleigh, E.A. and Wilson, G.G. (2014) Type I restriction enzymes and their relatives. Nucleic Acids Res., 42, 20–44.

36. De Ste Croix, M., Chen, K., Vacca, I., Manso, A., Johnston, C., Polard, P., Kwun, M., Bentley, S., Croucher, N. and Bayliss, C. (2019) Recombination of the phase-variable *spnIII* locus is independent of all known pneumococcal site-specific recombinases. J. Bacteriol., 201, e00233–00219.

37. Phillips, Z.N., Trappetti, C., Van Den Bergh, A., Martin, G., Calcutt, A., Ozberk, V., Guillon, P., Pandey, M., von Itzstein, M. and Swords, W.E. (2022) Pneumococcal phasevarions control multiple virulence traits, including vaccine candidate expression. *Microbiology Spectrum*, e00916–00922.

38. Oliver, M.B., Basu Roy, A., Kumar, R., Lefkowitz, E.J. and Swords, W.E. (2017) *Streptococcus pneumoniae* TIGR4 phase-locked opacity variants differ in virulence phenotypes. MSphere, 2, e00386–00317.

39. Atack, J.M., Weinert, L.A., Tucker, A.W., Husna, A.U., Wileman, T.M., F. Hadjirin, N., Hoa, N.T., Parkhill, J., Maskell, D.J. and Blackall, P.J. (2018) *Streptococcus suis* contains multiple phase-variable methyltransferases that show a discrete lineage distribution. Nucleic Acids Res., 46, 11466–11476.

40. Tram, G., Jen, F.E.-C., Phillips, Z.N., Timms, J., Husna, A.-U., Jennings, M.P., Blackall, P.J. and Atack, J.M. (2021) *Streptococcus suis* encodes multiple allelic variants of a phase-variable Type III DNA methyltransferase, ModS, that control distinct phasevarions. Msphere, 6, e00069–00021.

41. Turni, C. and Blackall, P. (2007) An evaluation of the *apxIVA* based PCR-REA method for differentiation of *Actinobacillus pleuropneumoniae*. Vet. Microbiol., 121, 163–169.

42. Bossé, J.T., Soares-Bazzolli, D.M., Li, Y., Wren, B.W., Tucker, A.W., Maskell, D.J., Rycroft, A.N., Langford, P.R. and Consortium, B.T. (2014) The generation of successive unmarked mutations and chromosomal insertion of heterologous genes in *Actinobacillus pleuropneumoniae* using natural transformation. PLoS One, 9, e111252.

43. Seemann, T. (2014) Prokka: rapid prokaryotic genome annotation. Bioinformatics, 30, 2068–2069.

44. Tonkin-Hill, G., MacAlasdair, N., Ruis, C., Weimann, A., Horesh, G., Lees, J.A., Gladstone, R.A., Lo, S., Beaudoin, C., Floto, R.A. et al. (2020) Producing polished prokaryotic pangenomes with the Panaroo pipeline. Genome biology, 21, 180.

45. Paradis, E. and Schliep, K. (2019) ape 5.0: an environment for modern phylogenetics and evolutionary analyses in R. Bioinformatics, 35, 526–528.

46. 46. Team, R.C. (2018) R: A language and environment for statistical computing. R Foundation for Statistical Computing. R Core, Vienna, Austria.

47. Letunic, I. and Bork, P. (2016) Interactive tree of life (iTOL) v3: an online tool for the display and annotation of phylogenetic and other trees. Nucleic Acids Res., 44, W242–W245.

48. Turni, C., Singh, R., Schembri, M. and Blackall, P. (2014) Evaluation of a multiplex PCR to identify and serotype *Actinobacillus pleuropneumoniae* serovars 1, 5, 7, 12 and 15. Lett. Appl. Microbiol., 59, 362–369.

49. Yee, S., Blackall, P. and Turni, C. (2018) Genetic diversity and toxin gene distribution among serovars of *Actinobacillus pleuropneumoniae* from Australian pigs. Aust. Vet. J., 96, 17–23.

50. Jansen, R., Briaire, J., Smith, H.E., Dom, P., Haesebrouck, F., Kamp, E.M., Gielkens, A. and Smits, M.A. (1995) Knockout mutants of *Actinobacillus pleuropneumoniae* serotype 1 that are devoid of RTX toxins do not activate or kill porcine neutrophils. Infect. Immun., 63, 27–37.

51. Bossé, J.T., Sinha, S., Schippers, T., Kroll, J.S., Redfield, R.J. and Langford, P.R. (2009) Natural competence in strains of *Actinobacillus pleuropneumoniae*. FEMS Microbiol. Lett., 298, 124–130.

52. Herriott, R.M., Meyer, E.M. and Vogt, M. (1970) Defined nongrowth media for stage II development of competence in *Haemophilus influenzae*. J Bacteriol, 101, 517–524.

53. Clark, T.A., Murray, I.A., Morgan, R.D., Kislyuk, A.O., Spittle, K.E., Boitano, M., Fomenkov, A., Roberts, R.J. and Korlach, J. (2011) Characterization of DNA methyltransferase specificities using single-molecule, real-time DNA sequencing. Nucleic Acids Res., 40, e29–e29.

54. Murray, I.A., Clark, T.A., Morgan, R.D., Boitano, M., Anton, B.P., Luong, K., Fomenkov, A., Turner, S.W., Korlach, J. and Roberts, R.J. (2012) The methylomes of six bacteria. Nucleic Acids Res., 40, 11450–11462.

55. Tourancheau, A., Mead, E.A., Zhang, X.-S. and Fang, G. (2021) Discovering multiple types of DNA methylation from bacteria and microbiome using nanopore sequencing. Nat. Methods, 18, 491–498.

56. Peak, I.R., Chen, A., Jen, F.E.-C., Jennings, C., Schulz, B.L., Saunders, N.J., Khan, A., Seifert, H.S. and Jennings, M.P. (2016) *Neisseria meningitidis* lacking the major porins PorA and PorB is viable and modulates apoptosis and the oxidative burst of neutrophils. Journal of proteome research, 15, 2356–2365.

57. Murphy, T.F., Dudas, K.C., Mylotte, J.M. and Apicella, M.A. (1983) A subtyping system for nontypable *Haemophilus influenzae* based on outer-membrane proteins. J. Infect. Dis., 147, 838–846.

58. Oakley, B.R., Kirsch, D.R. and Morris, N.R. (1980) A simplified ultrasensitive silver stain for detecting proteins in polyacrylamide gels. Anal. Biochem., 105, 361–363.

59. 59. CLSI (ed.) (2020) Performance standards for antimicrobial disk and dilution susceptibility tests for bacteria isolated from animals. 5th ed. Clinical and Laboratory Standards Institute Wayne, PA.

60. Labrie, J., Pelletier-Jacques, G., Deslandes, V., Ramjeet, M., Auger, E., Nash, J.H. and Jacques, M. (2010) Effects of growth conditions on biofilm formation by *Actinobacillus pleuropneumoniae*. Vet. Res., 41, 1–17.

61. Li, J., Li, J.W., Feng, Z., Wang, J., An, H., Liu, Y., Wang, Y., Wang, K., Zhang, X., Miao, Z. et al. (2016) Epigenetic switch driven by DNA inversions dictates phase variation in *Streptococcus pneumoniae*. PLoS Path., 12, e1005762.

62. Blakeway, L.V., Tan, A., Lappan, R., Ariff, A., Pickering, J.L., Peacock, C.S., Blyth, C.C., Kahler, C.M., Chang, B.J. and Lehmann, D. (2018) *Moraxella catarrhalis* restriction–modification systems are associated with phylogenetic lineage and disease. Genome Biology and Evolution, 10, 2932–2946.

63. Bossé, J.T., Li, Y., Walker, S., Atherton, T., Fernandez Crespo, R., Williamson, S.M., Rogers, J., Chaudhuri, R.R., Weinert, L.A. and Oshota, O. (2015) Identification of *dfrA14* in two distinct plasmids conferring trimethoprim resistance in *Actinobacillus pleuropneumoniae*. J. Antimicrob. Chemother., 70, 2217–2222.

64. De Ste Croix, M., Vacca, I., Kwun, M.J., Ralph, J.D., Bentley, S.D., Haigh, R., Croucher, N.J. and Oggioni, M.R. (2017) Phase-variable methylation and epigenetic regulation by type I restriction–modification systems. FEMS Microbiol. Rev., 41, S3–S15.

65. Zhou, L., Jones, S., Angen, Ø., Bossé, J., Nash, J., Frey, J., Zhou, R., Chen, H., Kroll, J. and Rycroft, A. (2008) Multiplex PCR that can distinguish between immunologically cross-reactive serovar 3, 6, and 8 *Actinobacillus pleuropneumoniae* strains. J. Clin. Microbiol., 46, 800–803.

66. Turni, C., Singh, R., Omaleki, L. and Blackall, P. (2013) *Actinobacillus pleuropneumoniae*–a diagnostic update. Australian Pig Science Association Meeting 2013

67. Murray, I.A., Luyten, Y.A., Fomenkov, A., Dai, N., Corrêa Jr, I.R., Farmerie, W.G., Clark, T.A., Korlach, J., Morgan, R.D. and Roberts, R.J. (2021) Structural and functional diversity among Type III restriction-modification systems that confer host DNA protection via methylation of the N4 atom of cytosine. Plos one, 16, e0253267.

68. Jen, F.E.-C., Seib, K.L. and Jennings, M.P. (2014) Phasevarions mediate epigenetic regulation of antimicrobial susceptibility in *Neisseria meningitidis*. Antimicrobial Agents and Chemotherapy, 58, 4219–4221.

69. 69. Åberg, A., Gideonsson, P., Vallström, A., Olofsson, A., Öhman, C., Rakhimova, L., Borén, T., Engstrand, L., Brännström, K. and Arnqvist, A. (2014) A repetitive DNA element regulates expression of the *Helicobacter pylori* sialic acid binding adhesin by a rheostat-like mechanism. PLoS Path., 10, e1004234.

70. Blyn, L.B., Braaten, B.A. and Low, D.A. (1990) Regulation of *pap* pilin phase variation by a mechanism involving differential dam methylation states. EMBO J., 9, 4045–4054.

71. Fox, K.L., Atack, J.M., Srikhanta, Y.N., Eckert, A., Novotny, L.A., Bakaletz, L.O. and Jennings, M.P. (2014) Selection for phase variation of LOS biosynthetic genes frequently occurs in progression of non-typeable *Haemophilus influenzae* infection from the nasopharynx to the middle ear of human patients. PLoS One, 9, e90505.

72. Poole, J., Foster, E., Chaloner, K., Hunt, J., Jennings, M.P., Bair, T., Knudtson, K., Christensen, E., Munson, R.S., Jr., Winokur, P.L. et al. (2013) Analysis of nontypeable *Haemophilus influenzae* phase-variable genes during experimental human nasopharyngeal colonization. Journal of Infectious Disease, 208, 720–727.

73. Blakeway, L.V., Tan, A., Jurcisek, J.A., Bakaletz, L.O., Atack, J.M., Peak, I.R. and Seib, K.L. (2019) The *Moraxella catarrhalis* phase-variable DNA methyltransferase ModM3 is an epigenetic regulator that affects bacterial survival in an *in vivo* model of otitis media. BMC Microbiol., 19, 1–14.

74. Gouil, Q. and Keniry, A. (2019) Latest techniques to study DNA methylation. Essays Biochem, 63, 639–648.

75. McIntyre, A.B.R., Alexander, N., Grigorev, K., Bezdan, D., Sichtig, H., Chiu, C.Y. and Mason, C.E. (2019) Single-molecule sequencing detection of N6-methyladenine in microbial reference materials. Nature communications, 10, 579.

76. Blakeway, L.V., Tan, A., Peak, I.R., Atack, J.M. and Seib, K.L. (2020) Proteome of a *Moraxella catarrhalis* strain under iron-restricted conditions. Microbiology Resource Announcements, 9, e00064–00020.

77. Brockman, K.L., Azzari, P.N., Branstool, M.T., Atack, J.M., Schulz, B.L., Jen, F.E.-C., Jennings, M.P. and Bakaletz, L.O. (2018) Epigenetic regulation alters biofilm architecture and composition in multiple clinical isolates of nontypeable *Haemophilus influenzae*. MBio, 9, e01682–01618.

78. Zhang, L., Luo, W., Xiong, R., Li, H., Yao, Z., Zhuo, W., Zou, G., Huang, Q. and Zhou, R. (2022) A combinatorial vaccine containing inactivated bacterin and subunits provides protection against *Actinobacillus pleuropneumoniae* infection in mice and pigs. Frontiers in veterinary science, 9.

79. Hölzen, P., Warnck, T., Hoy, S., Schlegel, K., Hennig-Pauka, I. and Gaumann, H. (2021) Comparison of protectivity and safety of two vaccines against *Actinobacillus pleuropneumoniae* in a field study. Agriculture, 11, 1143.

80. Shao, M., Wang, Y., Wang, C., Guo, Y., Peng, Y., Liu, J., Li, G., Liu, H. and Liu, S. (2010) Evaluation of multicomponent recombinant vaccines against *Actinobacillus pleuropneumoniae* in mice. Acta Veterinaria Scandinavica, 52, 1–8.

81. Byrd, W., Harmon, B.G. and Kadis, S. (1992) Protective efficacy of conjugate vaccines against experimental challenge with porcine *Actinobacillus pleuropneumoniae*. Vet. Immunol. Immunopathol., 34, 307–324.

82. Hathroubi, S., Loera-Muro, A., Guerrero-Barrera, A.L., Tremblay, Y.D. and Jacques, M. (2018) *Actinobacillus pleuropneumoniae* biofilms: role in pathogenicity and potential impact for vaccination development. Animal health research reviews, 19, 17–30.

83. Nahar, N., Turni, C., Tram, G., Blackall, P.J. and Atack, J.M. (2021) *Actinobacillus pleuropneumoniae*: the molecular determinants of virulence and pathogenesis. Adv. Microb. Physiol., 78, 179–216.

84. Yan, J. and Bassler, B.L. (2019) Surviving as a community: antibiotic tolerance and persistence in bacterial biofilms. Cell host & microbe, 26, 15–21.

85. Murphy, T.F., Bakaletz, L.O. and Smeesters, P.R. (2009) Microbial interactions in the respiratory tract. The Pediatric infectious disease journal, 28, S121–S126.

86. Novotny, L.A., Brockman, K.L., Mokrzan, E.M., Jurcisek, J.A. and Bakaletz, L.O. (2019) Biofilm biology and vaccine strategies for otitis media due to nontypeable *Haemophilus influenzae*. Journal of pediatric infectious diseases, 14, 069–078.

87. Perez-Riverol, Y., Csordas, A., Bai, J., Bernal-Llinares, M., Hewapathirana, S., Kundu, D.J., Inuganti, A., Griss, J., Mayer, G. and Eisenacher, M. (2019) The PRIDE database and related tools and resources in 2019: improving support for quantification data. Nucleic Acids Res., 47, D442–D450.

